# Nanopore sequencing and comparative genome analysis confirm lager-brewing yeasts originated from a single hybridization

**DOI:** 10.1101/603480

**Authors:** Alex N. Salazar, Arthur R. Gorter de Vries, Marcel van den Broek, Nick Brouwers, Pilar de la Torre Cortès, Niels G. A Kuijpers, Jean-Marc G. Daran, Thomas Abeel

**Affiliations:** Delft Bioinformatics Lab, Delft University of Technology, 2628 CD Delft, The Netherlands; Broad Institute of MIT and Harvard, Boston, MA 02142, USA; Department of Biotechnology, Delft University of Technology, Van der Maasweg 9, 2629 HZ Delft, The Netherlands; HEINEKEN Supply Chain B.V., Global Innovation and Research, Zoeterwoude, Netherlands

## Abstract

**Background:** The lager brewing yeast, *S. pastorianus*, is a hybrid between *S. cerevisiae* and *S. eubayanus* with extensive chromosome aneuploidy. *S. pastorianus* is subdivided into Group 1 and Group 2 strains, where Group 2 strains have higher copy number and a larger degree of heterozygosity for *S. cerevisiae* chromosomes. As a result, Group 2 strains were hypothesized to have emerged from a hybridization event distinct from Group 1 strains. Current genome assemblies of *S. pastorianus* strains are incomplete and highly fragmented, limiting our ability to investigate their evolutionary history.

**Results:** To fill this gap, we generated a chromosome-level genome assembly of the *S. pastorianus* strain CBS 1483 using MinION sequencing and analysed the newly assembled subtelomeric regions and chromosome heterozygosity. To analyse the evolutionary history of *S. pastorianus* strains, we developed Alpaca: a method to compute sequence similarity between genomes without assuming linear evolution. Alpaca revealed high similarities between the *S. cerevisiae* subgenomes of Group 1 and 2 strains, and marked differences from sequenced *S. cerevisiae strains*.

**Conclusions:** Our findings suggest that Group 1 and Group 2 strains originated from a single hybridization involving a heterozygous *S. cerevisiae* strain, followed by different evolutionary trajectories. The clear differences between both groups may originate from a severe population bottleneck caused by the isolation of the first pure cultures. Alpaca provides a computationally inexpensive method to analyse evolutionary relationships while considering non-linear evolution such as horizontal gene transfer and sexual reproduction, providing a complementary viewpoint beyond traditional phylogenetic approaches.

## Background

The lager-brewing yeast *Saccharomyces pastorianus* is an interspecies hybrid between *S. cerevisiae* and *S. eubayanus*. Lager brewing emerged in the late middle ages and was carried out during winter months at temperatures between 8 and 15 °C, followed by a prolonged maturation period referred to as lagering (1, 2). While *S. cerevisiae* is a well-studied species frequently used in biotechnological processes (3), *S. eubayanus* was only discovered in 2011 and has thus far only been isolated from the wild (4). Therefore, the ancestral *S. pastorianus* hybrid likely emerged from a spontaneous hybridization between an ale brewing *S. cerevisiae* yeast and a wild *S. eubayanus* contaminant, and took over lager brewing due to increased fitness under these conditions (4–6). Indeed, laboratory-made *S. cerevisiae* × S. *eubayanus* hybrids demonstrated hybrid vigour by combining the fermentative capacity and sugar utilisation of *S. cerevisiae* and the ability to grow at lower temperatures of *S. eubayanus* (7, 8).

The genomes of *S. pastorianus* strains are highly aneuploid, containing 0 to 5 copies of each chromosome (5, 9-13). Between 45 and 79 individual chromosomes were found in individual *S. pastorianus* genomes, compared to a normal complement of 32 chromosomes in euploid *Saccharomyces* hybrids. The degree of aneuploidy of *S. pastorianus* is exceptional in the *Saccharomyces* genera, and likely evolved during its domestication in the brewing environment (9). Nevertheless, two groups can be distinguished based on their genome organisation: Group 1 strains, which have approximately haploid *S. cerevisiae* and diploid *S. eubayanus* chromosome complements; and Group 2 strains, which have approximately diploid to tetraploid *S. cerevisiae* and diploid *S. eubayanus* chromosome complements (5, 10, 11, 14).

Group 1 and Group 2 strains in *S. pastorianus* were initially thought to have originated from two different hybridization events. Some lager-specific genes from Group 2 strains are absent in Group 1 strains, and the subtelomeric regions of Group 1 and Group 2 strains differ substantially (15, 16). Based on these differences, Group 1 and Group 2 strains were hypothesized to have emerged from different independent hybridization events, involving a haploid *S. cerevisiae* for Group 1 strains and a higher ploidy *S. cerevisiae* strain for Group 2 strains (5, 17). Indeed, crosses between *S. cerevisiae* and *S. eubayanus* strains with varying ploidies could be made in the laboratory, all of which performed well in the lager brewing process (18). Comparative genome analysis between Group 1 and Group 2 strains revealed that there were more synonymous nucleotide differences in the *S. cerevisiae* subgenome than in the *S. eubayanus* subgenome (19). As accumulation of synonymous mutations was presumed to equally affect both genomes, the authors hypothesized that Group 1 and 2 strains originated from two hybridizations, with a similar *S. eubayanus* parent and different *S. cerevisiae* parents.

More recent studies now support that Group 1 and Group 2 strains originated from the same hybridization event. Identical recombinations between the *S. cerevisiae* and *S. eubayanus* subgenomes were found at the *ZUO1*, *MAT*, *HSP82* and *XRN1*/*KEM1* loci in all analysed *S. pastorianus* strains (11, 13, 14), which did not emerge when such hybrids were evolved under laboratory conditions (20). These conserved recombinations indicate that all *S. pastorianus* strains share a common *S. cerevisiae* × *S. eubayanus* hybrid ancestor, and that the differences between Group 1 and Group 2 strains emerged subsequently. Sequence analysis of ten *S. pastorianus* genomes revealed that the *S. cerevisiae* sub-genome in Group 1 strains is relatively homozygous, while Group 2 strains possess heterozygous sub-regions (11). Moreover, heterozygous nucleotide stretches in Group 2 strains were composed of sequences highly similar to Group 1 genomes and of sequences from a different *S. cerevisiae* genome with a 0.5% lower sequence identity. As a result, the authors formulated two hypotheses to explain the emergence of Group 1 and Group 2 strains from a shared ancestral hybrid: (i) the ancestral hybrid had a heterozygous *S. cerevisiae* sub-genome, and Group 1 strains underwent a massive reduction of the *S. cerevisiae* genome content while Group 2 did not, or (ii) the ancestral hybrid had a homozygous Group 1-like genome and Group 2 strains were formed by a subsequent hybridization event of such a Group 1-like strain with another *S. cerevisiae* strain, resulting in a mixed *S. cerevisiae* genome content in Group 2 strains.

Since the exact *S. cerevisiae* and *S. eubayanus* ancestors of *S. pastorianus* are not available, the evolutionary history of *S. pastorianus* has so far been based on the sequence analysis using available *S. cerevisiae* and *S. eubayanus* reference genomes (5, 11). However, these reference genomes are not necessarily representative of the original parental genomes of *S. pastorianus*. Although *S. pastorianus* genomes are available, they were sequenced with short-read sequencing technology (10–13) preventing assembly of large repetitive stretches of several thousand base pairs, such as TY-elements or paralogous genes often found in *Saccharomyces* genomes (21). The resulting *S. pastorianus* genomes assemblies are thus incomplete and fragmented into several hundred or thousand contigs (10–13).

Single-molecule sequencing technologies can output reads of several thousand base pairs and span entire repetitive regions, enabling near complete chromosome-level genome assemblies of *Saccharomyces* yeasts (22–27). In addition to the lesser fragmentation, the assembly of regions containing repetitive sequences reveals large numbers of previously unassembled open reading frames, particularly in the sub-telomeric regions of chromosomes (24, 25, 27). Sub-telomeric regions are relatively unstable (28), and therefore contain much of the genetic diversity between different strains (29, 30). In *S. pastorianus*, notable differences were found between the sub-telomeric regions of Group 1 and Group 2 strains (15, 16), which could be used to understand their origin. Moreover, repetitive regions are enriched for genes with functions determining the cell’s interaction with its environment, such as nutrient uptake, sugar utilization, inhibitor tolerance and flocculation (31–34). As a result, the completeness of sub-telomeric regions is critical for understanding genetic variation and evolutionary relationships between strains, as well as for understanding their performance in industrial applications (24, 29, 30).

Here, we used nanopore sequencing to obtain a chromosome-level assembly of the Group 2 *S. pastorianus* strain CBS 1483 and analysed the importance of new-found sequences relative to previous genome assemblies, with particular focus on industrially-relevant subtelomeric gene families. As the CBS 1483 genome contains multiple non-identical copies for many chromosomes, we analysed structural and sequence-level heterozygosity using short- and long-read data. Moreover, we developed a method to investigate the evolutionary origin of *S. pastorianus* by evaluating the genome similarity of several Group 1 and Group 2 *S. pastorianus* strains relative to a large dataset of *S. cerevisiae* and *S. eubayanus* genomes, including an isolate of the Heineken A-yeast^®^ lineage which was isolated by dr. Elion in 1886 and is still used in beer production today.

## Results

### Near-complete haploid assembly of CBS 1483

We obtained 3.3 Gbp of whole genome sequencing data of the *Saccharomyces pastorianus* strain CBS 1483 using 4 flow cells on Oxford Nanopore Technology’s MinION platform. Based on a genome size of 46 Mbp accounting for all chromosome copy numbers, the combined coverage was 72x with an average read length of 7 Kbp (Figure S1). We assembled the reads using Canu (35) and performed manual curation involving circularization of the mitochondrial DNA, scaffolding of *Sc*XII (chromosome XII of the *S. cerevisiae* sub-genome) and resolution of assembly problems due to inter- and intra-chromosomal structural heterozygosity in *Sc*I and *Sc*XIV (Figure 1). Assembly errors were corrected with Pilon (36) using paired-end Illumina reads with 159x coverage. We obtained a final assembly of 29 chromosome contigs, 2 chromosome scaffolds, and the complete mitochondrial contig leading to a total size of 23.0 Mbp (Figure 2 and Table 1). The assembly was remarkably complete: of the 31 chromosomes (in CBS 1483 *Sc*III and *Se*III recombined into a chimeric *Se*III-*Sc*III chromosome(10), 29 were in single contigs; 21 of the chromosomes contained both telomere caps; 8 contained one of the caps; and 2 were missing both caps. Some chromosomes contain sequence from both parental sub-genomes due to recombinations; those chromosomes were named *Se*III-*Sc*III, *Se*VII-*Sc*VII, *Sc*X-*Se*X, *Se*X-*Sc*X and *Se*XIII-ScXIII, in accordance with previous nomenclature (10). Annotation of the assembly resulted in the identification of 10,632 genes (Additional File 1A). We determined chromosome copy number based on coverage analysis of short-read alignments to the genome assembly of CBS 1483 (Figure 2 and S2).

**Figure 1.**
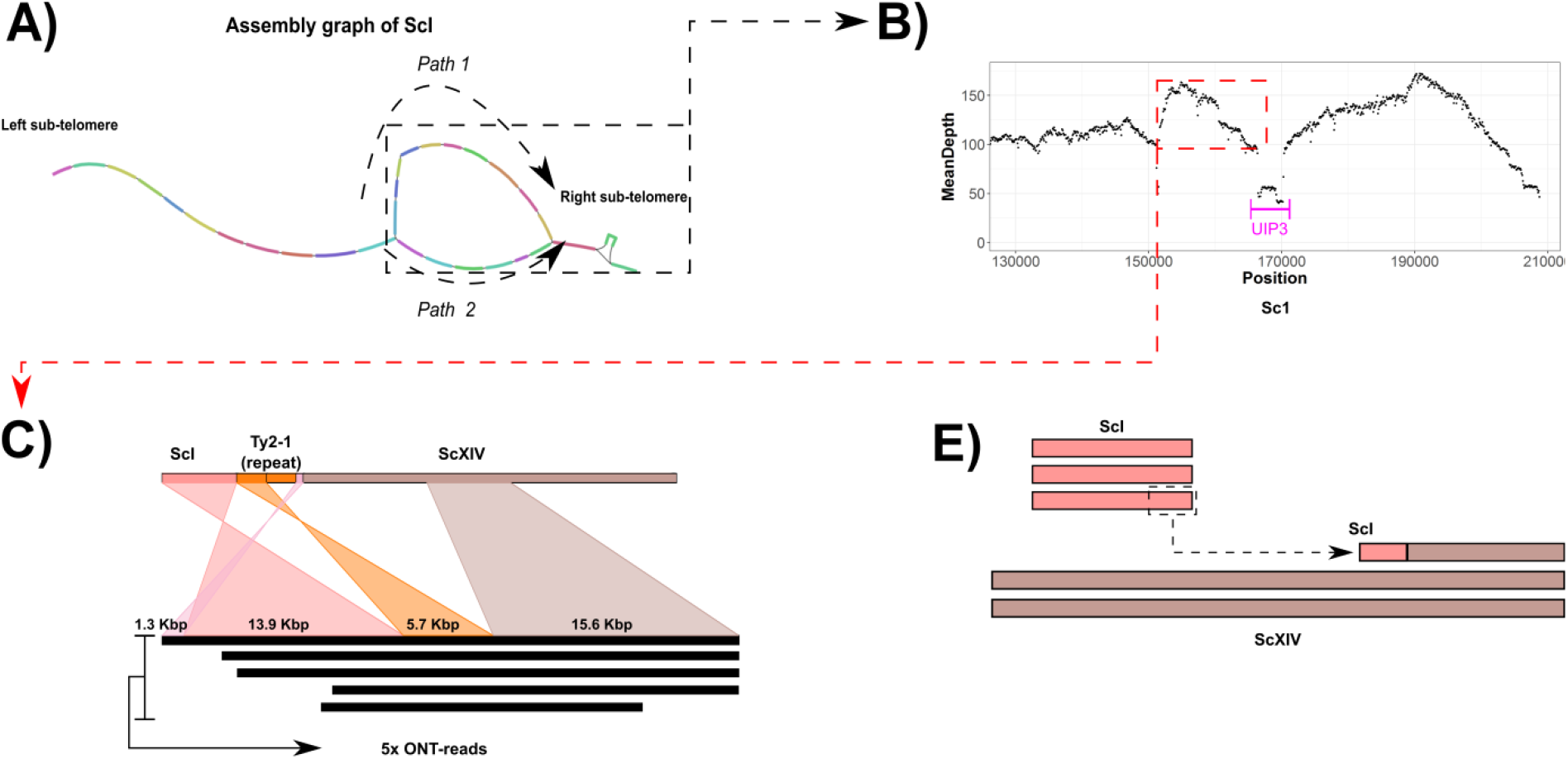
Structural heterozygosity within multiple copies of the *S. cerevisiae* chromosome I of CBS 1483. (A) Layout of *S. cerevisiae* chromosome I in the assembly graph. Paths 1 and 2 represent alternative contigs in the right-end of the chromosome—the gene *UIP3* is deleted in path 2. (B) Sequencing coverage of ONT read-alignments of CBS 1483 in the right-end of chromosome I after joining path 1 and discarding path 2. The location of the *UIP3* gene is indicated. (C) Alignment overview of five raw ONT reads supporting the introgression of a ∼14 Kbp in chromosome I (salmon colour) to a region at the right-end of chromosome XIV (brown colour) in the *S. cerevisiae* sub-genome. The additional alignments (pink and orange) are alignments to computationally confirmed Ty-2 repetitive elements. (D) Schematic representation of the two chromosome architectures of *S. cerevisiae* chromosome XIV (brown colour) due to translocation of an additional copy of the right arm of chromosome I (salmon colour) to the left arm of chromosome XIV.

**Figure 2.**
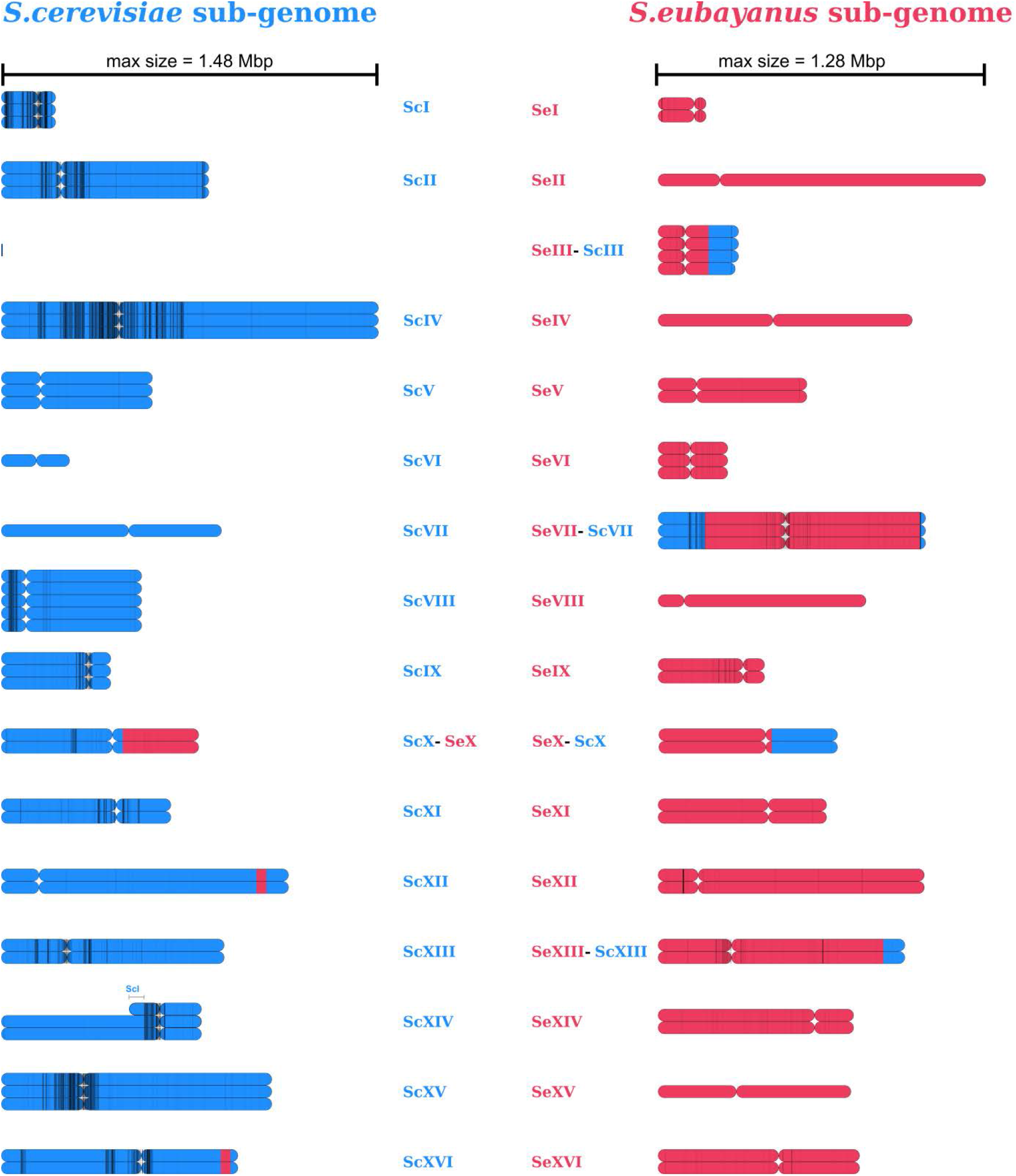
Overview of Nanopore-only *de novo* genome assembly of the *S. pastorianus* strain, CBS 1483. For each chromosome, all copies are represented. Genomic material originating from *S. cerevisiae* (blue) and from *S. eubayanus* (red) are shown, and the position of the centromere is indicated. Heterozygous SNP calls are represented as vertical, black lines and are drawn with transparency to depict the density of SNP calls in a given region. Underlying chromosome copy number data and the list of heterozygous SNPs is available in Figure S2 and Additional File 1F.

**Table 1.** Length and gaps of each assembled chromosome of the *S. cerevisiae* and *S. eubayanus* subgenome in the *de novo* assembly of Group 2 *S. pastorianus* strain CBS 1483. The mitochondrial DNA assembly is also shown.

| *S. cerevisiae* sub-genome |  |  | *S. eubayanus* sub-genome |  |  |
| --- | --- | --- | --- | --- | --- |
| Contig/Scaffold | Size | Gaps | Contig/Scaffold | Size | Gaps |
| ScI | 208794 | 0 | SeI | 183365 | 0 |
| ScII | 812290 | 0 | SeII | 1284912 | 0 |
| ScIII | 0 | 0 | SeIII | 311639 | 0 |
| ScIV | 1480484 | 0 | SeIV | 995872 | 0 |
| ScV | 590259 | 0 | SeV | 580717 | 0 |
| ScVI | 263951 | 0 | SeVI | 268897 | 0 |
| ScVII | 862436 | 0 | SeVII | 1048199 | 0 |
| ScVIII | 547874 | 0 | SeVIII | 813607 | 0 |
| ScIX | 426203 | 0 | SeIX | 413986 | 0 |
| ScX | 772632 | 0 | SeX | 698708 | 0 |
| ScXI | 662864 | 0 | SeXI | 658371 | 0 |
| ScXII | 1128411 | 2 | SeXII | 1043408 | 0 |
| ScXIII | 872991 | 0 | SeXIII | 966749 | 0 |
| ScXIV | 783474 | 0 | SeXIV | 765784 | 1 |
| ScXV | 1060500 | 0 | SeXV | 754183 | 0 |
| Sc XVI | 926828 | 0 | SeXVI | 788293 | 0 |
| Unplaced | 36198 | 0 | Mitochondria | 68765 | 0 |

### Comparison between ONT and Illumina assemblies

In order to compare our novel nanopore assembly of CBS 1483 to the previous assembly generated using short-read data, we aligned contigs of CBS 1483 from van den Broek *et al.* (10) to our current ONT-assembly, revealing a total 1.06 Mbp of added sequence. The added sequence overlapped with 323 ORFs (Additional File 1B). Conversely, aligning the nanopore assembly to the van den Broek *et al.* 2017 assembly revealed that only 14.9 Kbp of sequence was lost, affecting 15 ORFs (Additional File 1C). Gene ontology analysis of the added genes showed enrichment of several biological processes, functions, and components such as flocculation (P-value = 7.44×10^−3^) as well as transporter activity for several sugars including mannose, fructose and glucose (P-value ≤ 1.5×10^−5^) (Additional File 1D). Among the added genes were various members of subtelomeric gene families such as the *FLO*, *SUC*, *MAL*, *HXT* and *IMA* genes (Additional file 1E). Due to their role in the brewing-relevant traits such as carbohydrate utilization and flocculation, the complete assembly of subtelomeric gene families is crucial to capture different gene versions and copy number effects.

The assembly of CBS 1483 contained 9 *MAL* transporters, which encode for the ability to import maltose and maltotriose (37–39), constituting 85% of fermentable sugar in brewer’s wort (40). The *S. cerevisiae* subgenome harboured *ScMAL31* on *Sc*II, *ScMAL11* on *Sc*VII and on *Se*VII-*Sc*VII, and *ScMAL41* on *Sc*XI (Additional File 1B and 1E). However, the *ScMAL11* gene, also referred to as *AGT1*, was truncated, and there was no *ScMAL21* gene due to the complete absence of *Sc*III, as reported previously (10, 12). In the *S. eubayanus* subgenome, *MAL31*-type transporter genes were found in *Se*II, *Se*V, and *Se*XIII-ScXIII, corresponding to the location of the *S. eubayanus* transporter genes *SeMALT1*, *SeMALT2* and *SeMALT3*, respectively (25). In addition, a *MAL11*-like transporter was found on *Se*XV. Consistently with previous reports, no *MTY1*-like maltotriose transporter was found in CBS 1483 (10). Due to the absence of *MTY1* and the truncation of *ScMAL11*, maltotriose utilisation is likely to rely on the *SeMAL11* transporter in CBS 1483. Indeed, a *MAL11*-like transporter was recently shown to confer maltotriose utilisation in an *S. eubayanus* isolate from North Carolina (41).

The assembly also contained 14 *FLO* genes encoding flocculins which cause cell mass sedimentation upon completion of sugar consumption (34, 42, 43). The heavy flocculation of *S. pastorianus* cells simplifies biomass separation at the end of the brewing process, and resulted in their designation as bottom-fermenting yeast (44). Flocculation is mediated by flocculins: lectin-like cell wall proteins which effect cell-to-cell adhesion. In CBS 1483, we identified 12 flocculin genes, in addition to two *FLO8* transcriptional activators of flocculins (Additional File 1E). Flocculation intensity has been correlated to the length of flocculin genes (45–47). Specifically, increased length and number of tandem repeats within the *FLO* genes caused increased flocculation (47, 48). We therefore analysed tandem repeats in *S. cerevisiae*, *S. eubayanus* and *S. pastorianus* genomes and found that most *FLO* genes contain a distinct repeat pattern: two distinct, adjacent sequences each with variable copy number (Table 2). The repeats in *FLO1, FLO5, and FLO9* of the *S. cerevisiae* strain S288C have the same repeats of 135 bp and 15 bp; while repeats are of 189 bp and 15 bp for *FLO10* and of 132 bp and 45 bp for *FLO11*. The same repeat structures can be found in the *S. eubayanus* strain CBS 12357 as *FLO1, FLO5, and FLO9* contain repeats of 156 and 30 bp; although we were unable to find clear repeat patterns for *FLO10* and *FLO11* in this genome. In *S. pastorianus* CBS 1483, the repeat lengths of *FLO* genes corresponded to the subgenome they were localized in (Table 2). Compared to the non-flocculent S288C and CBS 12357 strains, *FLO* genes were systematically shorter in CBS 1483, contrasting with available theory (42–50). The intense flocculation phenotype of *S. pastorianus* was previously attributed to a gene referred to as *LgFLO1* (49, 51, 52). However, alignment of previously published partial and complete *LgFLO1* sequences did not confirm the presence of a similar ORF in CBS 1483. Moreover, the annotated *FLO* genes had higher identity with *S. eubayanus* and *S. cerevisiae FLO* genes, than with *LgFLO1.* Therefore, flocculation is likely to rely on one or several of the identified *FLO* genes from *S. cerevisiae* or *S. eubayanus* subgenomes (Table 2).

**Table 2.**
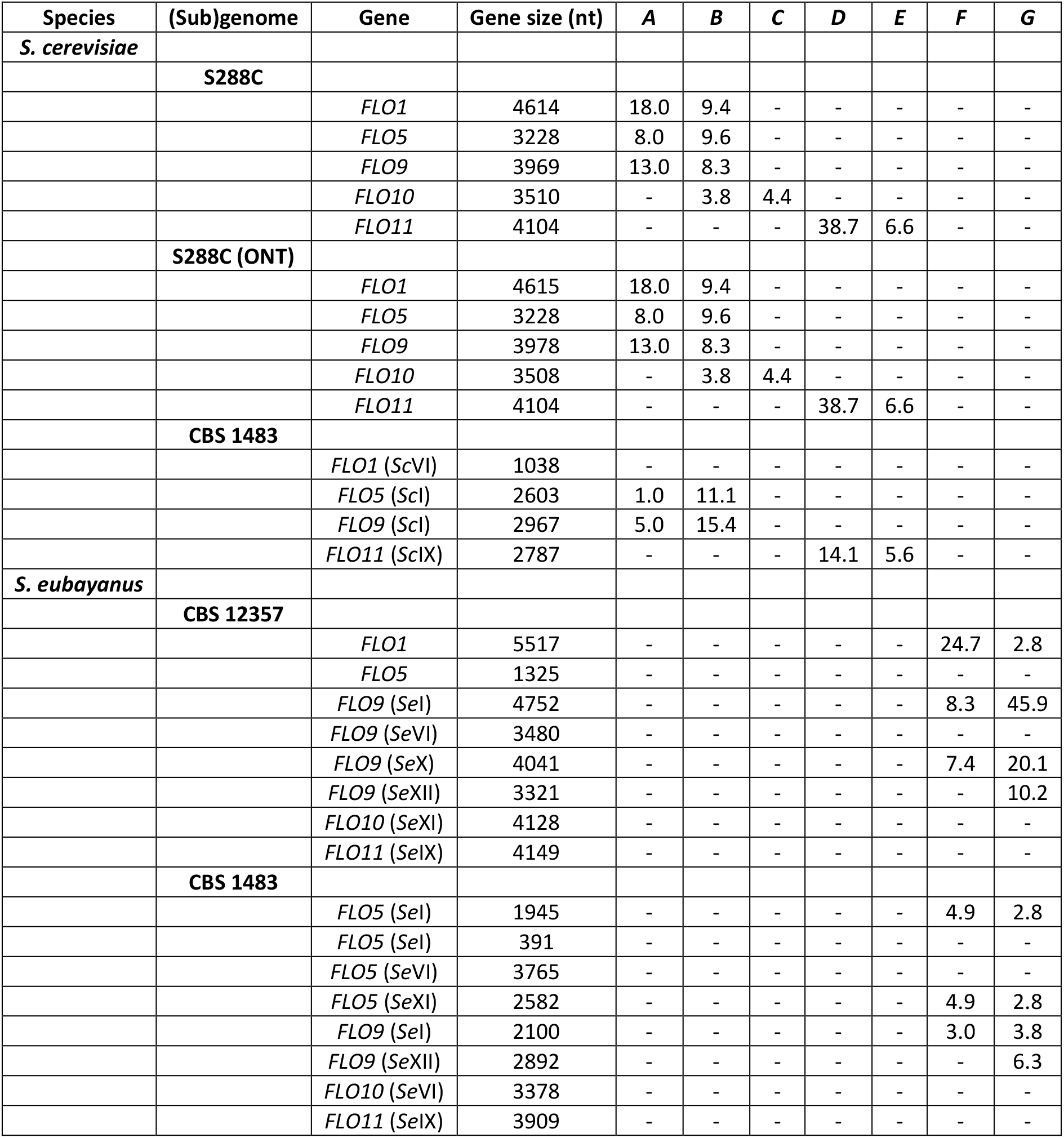
Tandem repeat analysis in *FLO* genes. We found seven repeat sequences when analysing flocculation genes *FLO1, FLO5, FLO9, FLO10, and FLO11* in *S. cerevisiae* (S288C) and *S. eubayanus* (CBS 12357) genomes. These sequences are referred to as sequence A (135 nt), B (15 nt), C (189 nt), D (45 nt), E (132 nt), F (156 nt), and G (30 nt). We used these sequences to analyse the copy numbers of each repeat within all *FLO* genes in our ONT-only assembly of CBS 1483 using the ONT-only S288C assembly as a control. Their respective copy numbers are shown below. Repeat sequences are indicated in Additional File 1H.

### Sequence heterogeneity in CBS 1483

As other Group 2 *S. pastorianus* strains, CBS 1483 displays heterozygosity between different copies of its *S. cerevisiae* subgenome (11). We therefore systematically identified heterozygous nucleotides in its genome and investigated the ORFs with allelic variation. Using 156x coverage of paired-end Illumina library of CBS 1483, we found a total of 6,367 heterozygous SNPs across the genome (Additional File 1F). Although the heterozygous SNPs are present across the whole genome, they affect primarily the *S. cerevisiae* sub-genome, with the majority clustered around centromeres (Figure 2). Of these positions, 58% were located within ORFs, resulting in 896 ORFs with allelic variation consisting of 1 to 30 heterozygous nucleotides. A total of 685 ORFs showed heterozygosity which would result in amino acid sequence changes, including 16 premature stop codons, 4 lost stop codons and 1566 amino acid substitutions (Additional File 1F). Gene ontology analysis of the ORFs affected by heterozygous calls revealed no significant enrichment in processes, functions of compartments. However, it should be noted that several industrially-relevant genes encoded more than one protein version, such as: the *BDH1* and *BDH2* genes, encoding butane-diol dehydrogenases involved in reduction of the off flavour compound diacetyl (*53*), the *FLO5* and *FLO9* genes encoding flocculins (50), and the *OAF1* gene encoding a regulator of ethyl-ester production pathway (54).

### Structural heterogeneity in CBS 1483 chromosomes

We investigated whether information about structural heterogeneity between chromosome copies could be recovered despite the fact that current assembly algorithms reduce genome assemblies to consensus sequences. Information about structural and sequence variation between different chromosome haplotypes is not captured by consensus assemblies. However, raw read data contains information for each chromosome copy. To identify structural heterogeneity, we identified ORFs whose predicted copy number deviated from that of the surrounding region in the chromosome based on read coverage analysis (Figure S3). We found 213 ORFs with deviating copy number (Additional File 1G). While no enrichment was found by gene ontology analysis, many of these ORFs are located in subtelomeric regions (29). Nevertheless, a few regions contained adjacent ORFs with deviating copy number, indicating larger structural variation between chromosome copies. For example, 21 consecutive ORFs in the right-end of the *Sc*XV appear to have been deleted in 2 of the 3 chromosome copies (Figure S3). *UIP3*, one of the genes with deviating copy number, was located on the right arm of chromosome *Sc*I. This region was previously identified as having an additional copy in CBS 1483, although it could not be localized based on short read data (10). The assembly graph showed two possible structures for *Sc*I, which were collapsed into a single contig in the final assembly (Figure 1A). Sequence alignment, gene annotations and sequencing coverage indicated two versions of the *Sc*I contigs: one with and one without the gene *UIP3* (Figure 1B). Sequence alignments of raw-ONT reads revealed five reads (from 20.6 to 36.7 Kbp) linking the right arm of *Sc*I to the left arm of *Sc*XIV at position ∼561 Kbp (Figure 1C). This location corresponded to a Ty-2 repetitive element; known to mediate recombination within *Saccharomyces* genomes (21). In addition to the increased coverage of the right arm of *Sc*I, the left arm of *Sc*XIV showed decreased sequencing coverage up until the ∼561 Kbp position. Together, these results suggest that the left arm of one copy of *Sc*XIV was replaced with an additional copy of the right arm of *Sc*I (Figure 1D). As no reads covered both the recombination locus and the *UIP3* locus, it remained unclear if *UIP3* is present in the *Sc*I copy translocated to chromosome *Sc*XIV. The resolution of two alternative chromosome architectures of *Sc*I and *Sc*XIV illustrates the ability of long-read alignment to resolve structural heterozygosity.

### Differences between Group 1 and 2 genomes do not result from separate ancestry

*S. pastorianus* strains can be subdivided into two separate groups—termed Group 1 and Group 2—based on both phenotypic (55) and genomic features (5, 11). However, the ancestral origin of each group remains unclear. The two groups may have emerged by independent hybridization events (19). Alternatively, Group 1 and Group 2 strains may originate from the same hybridization event, but Group 2 strains later hybridized with a different *S. cerevisiae* strain (11). In both cases, analysis of the provenance of genomic material from Group 1 and Group 2 genomes could confirm the existence of separate hybridization events if different ancestries are identified. Pan-genomic analysis of *S. cerevisiae* strains indicated that their evolution was largely non-linear, involving frequent horizontal gene transfer and sexual backcrossing events (56). Especially if the evolutionary ancestry of *S. pastorianus* involves admixture of different *S. cerevisiae* genomes (11), approaches considering only linear evolution such as phylogenetic trees are insufficient (57). Complex, non-linear evolutionary relationships could be addressed with network approaches (58). However, such algorithms are not yet fully mature and would involve extreme computational challenges (59, 60).

Therefore, we developed Alpaca: a simple and computationally inexpensive method to investigate complex non-linear ancestry via comparison of sequencing datasets (61). Alpaca is based on short-read alignment of a collection of strains to a partitioned reference genome, in which the similarity of each partition to the collection of strains is independently computed using k-mer sets (61). Reducing the alignments in each partition to k-mer sets prior to similarity analysis is computationally inexpensive. Phylogenetic relationships are also not recalculated, but simply inferred from previously available information on the population structure of the collection of strains (61). The partitioning of the reference genome enables the identification of strains with high similarity to different regions of the genome, enabling the identification of ancestry resulting from non-linear evolution. Moreover, since similarity analysis is based on read data, heterozygosity is taken into account.

We used Alpaca to identify the most similar lineages for all non-overlapping 2 Kbp sub-regions in the genome of the Group 2 *S. pastorianus* strain CBS 1483 using a reference dataset of 157 *S. cerevisiae* strains (62) and 29 *S. eubayanus* strains (63). We inferred population structures for both reference datasets by using previously defined lineages of each strain along with hierarchical clustering based on genome similarity using MASH (64). For the *S. eubayanus* subgenome, almost all sub-regions of CBS 1483 were most similar to strains from the Patagonia B – Holartic lineage (63) (Figure 3). In fact, 68% of all sub-regions were most similar to the Tibetan isolate CDFM21L.1 (65) and 27% to two highly-related North-American isolates (Figure 4), indicating a monophyletic ancestry of the *S. eubayanus* genome. Analysis of *S. pastorianus* strains CBS 2156 and WS 34/70 (Group2), and of CBS 1503, CBS 1513 and CBS 1538 (Group 1), indicated identical ancestry of their *S. eubayanus* subgenomes (Figure 4). Overall, we did not discern differences in the *S. eubayanus* subgenomes of *S. pastorianus* strains, which seem to descend from a strain of the Patagonia B – Holartic lineage and which is most closely related to the Tibetan isolate CDFM21L.1.

**Figure 3.**
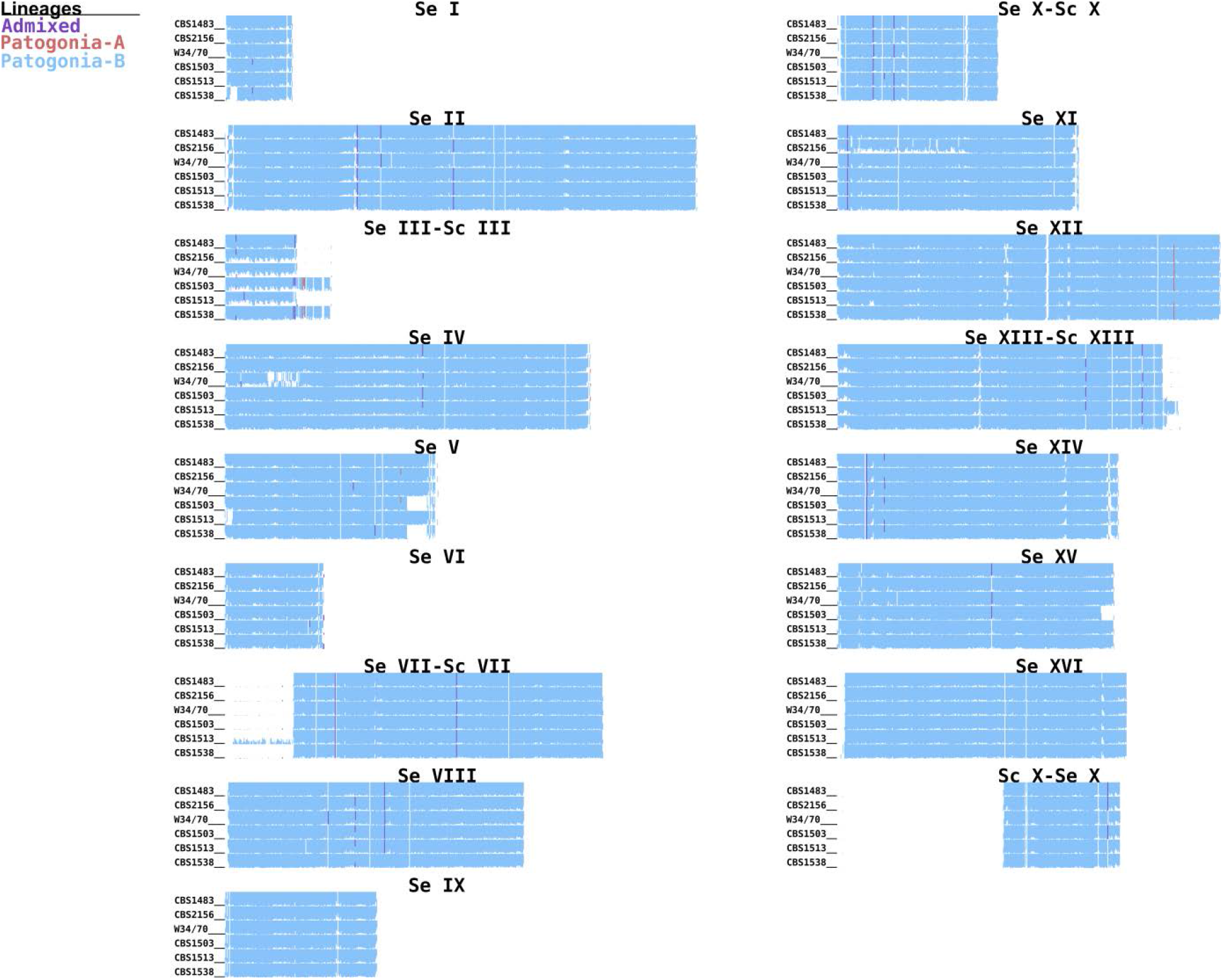
Similarity profiles of the *S. eubayanus* (sub-)genomes of Group 1 and 2 *S. pastorianus* strains, as determined using Alpaca. Each *S. eubayanus* chromosome of the CBS 1483 assembly was partitioned in non-overlapping sub-regions of 2 Kbp. The colors represent the most similar lineages based on k-mer similarity of 29 *S. eubayanus* strains from Peris *et al* (*63*): admixed (purple), Patagonia-A (red), Patagonia-B (blue). Similarity patterns are shown for the Group 2 strains CBS 1483, CBS 2156 and WS34/70 and the Group 1 strains CBS 1503, CBS 1513 and CBS 1538.

**Figure 4.**
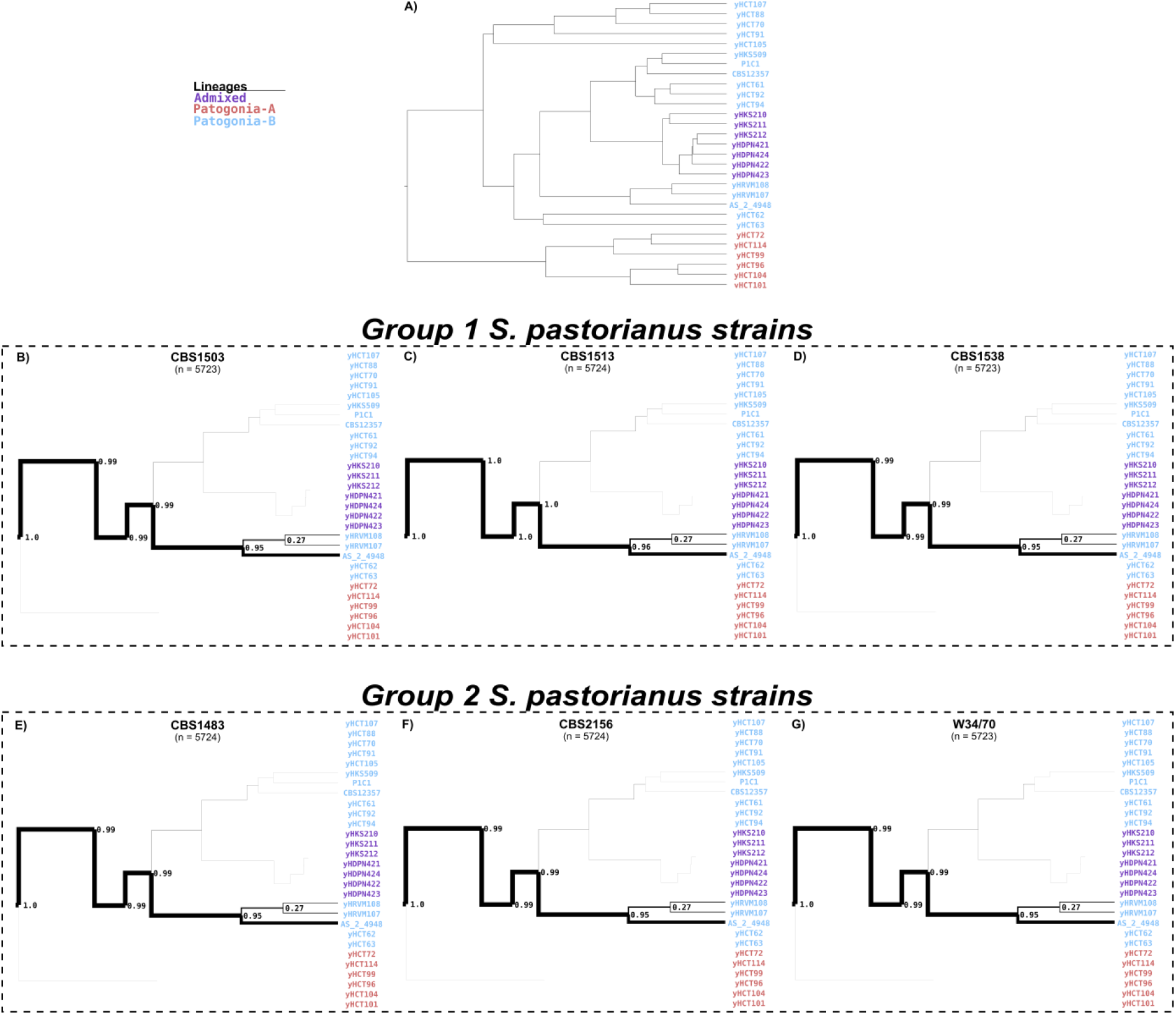
Tree-tracing of the genome-scale similarity across the *S. eubayanus* (sub-)genomes of Group 1 and 2 *S. pastorianus* strains, as determined using Alpaca. The frequency at which a genome from the reference data set of 29 *S. eubayanus* genomes from Peris *et al* (63) was identified as most similar for a sub-region of the CBS 1483 genome is depicted. The reference dataset is represented as a population tree, upon which only lineages with similarity are indicated with a thickness proportional to the frequency at which they were found as most similar (‘N’ being the total sum of the number of times all samples appeared as top-scoring). The complete reference population tree (A), the genomes of Group 1 strains CBS 1503, CBS 1513 and CBS 1538 (B-D) and for the genomes of Group 2 strains CBS 1483, CBS 2156 and WS34/70 (E-G) are shown.

In contrast, for the *S. cerevisiae* sub-genome of CBS 1483, the most similar *S. cerevisiae* strains varied across the sub-regions of every chromosome (Figure 5). No strain of the reference dataset was most similar for more than 5% of sub-regions, suggesting a high degree of admixture (Figure 6). However, 60% of sub-regions were most similar to the Beer 1 lineage, 12% were most similar to the Wine lineage and 10% to the Beer 2 lineage (62). In order to determine Alpaca’s ability to differentiate genomes with different admixed ancestries, we analysed the genomes of 8 *S. cerevisiae* strains: six ale-brewing strains and the laboratory strains CEN.PK113-7D and S288C. The strains CBS 7539, CBS 1463 and A81062 were identified as similar to the Beer 2 lineage, CBS 1171 and CBS 6308 as similar to the Beer 1 lineage, CBS 1487 as similar to the Wine lineage, and CEN.PK113-7D and S288C as similar to the mosaic laboratory strains (Figure 5). In addition, the distribution of similarity over the *S. cerevisiae* population tree differed per strain (Figure 6 and S4). While no single strain was most similar for more than 8% of the sub-regions for CBS 1487 and CBS 6308, for CBS 7539 67% of sub-regions were most similar to the strain beer002. As both beer002 and CBS 7539 are annotated as Bulgarian beer yeast (56, 62), this similarity likely reflects common origin. The different similarity profiles of all *S. cerevisiae* strains indicate that Alpaca can differentiate different ancestry by placement of genetic material within the *S. cerevisiae* population tree, whether a genome has a linear monophyletic origin or a non-linear polyphyletic origin.

**Figure 5.**
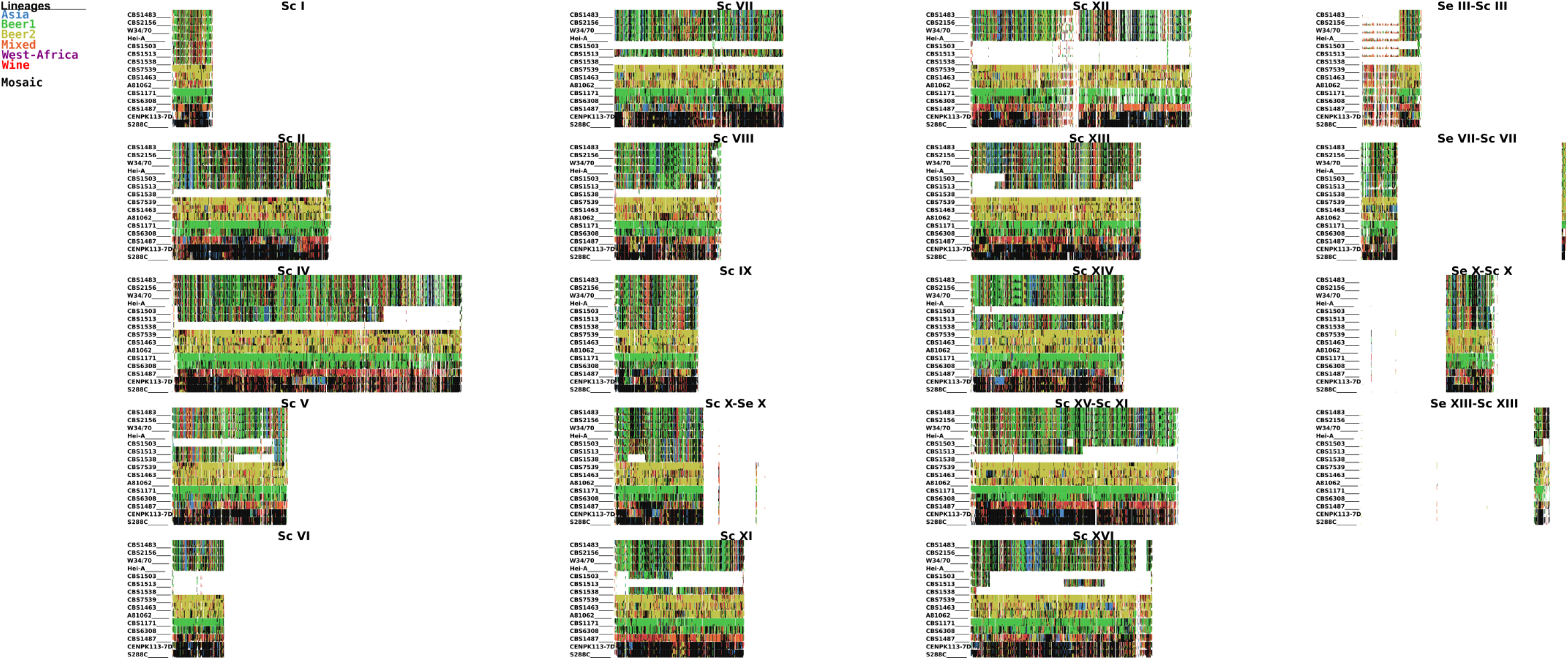
Similarity profiles of the *S. cerevisiae* (sub-)genomes of various *Saccharomyces* strains, as determined using Alpaca. Each *S. cerevisiae* chromosome of the CBS 1483 assembly was partitioned in non-overlapping sub-regions of 2 Kbp. The colors represent the most similar lineages based on k-mer similarity of 157 *S. cerevisiae* strains from Gallone *et al* (*62*): Asia (blue), Beer1 (green), Beer2, (gold), Mixed (orange), West-Africa (purple), Wine (red). Mosaic strains are shown in black and ambiguous or low-similarity sub-regions in white. Similarity patterns are shown for the Group 2 *S. pastorianus* strains CBS 1483, CBS 2156, WS34/70 and Hei-A, for the Group 1 *S. pastorianus* strains CBS 1503, CBS 1513 and CBS 1538, for *S. cerevisiae* ale-brewing strains CBS 7539, CBS 1463, A81062, CBS 1171, CBS 6308 and CBS 1483, and for *S. cerevisiae* laboratory strains CEN.PK113-7D and S288C.

**Figure 6.**
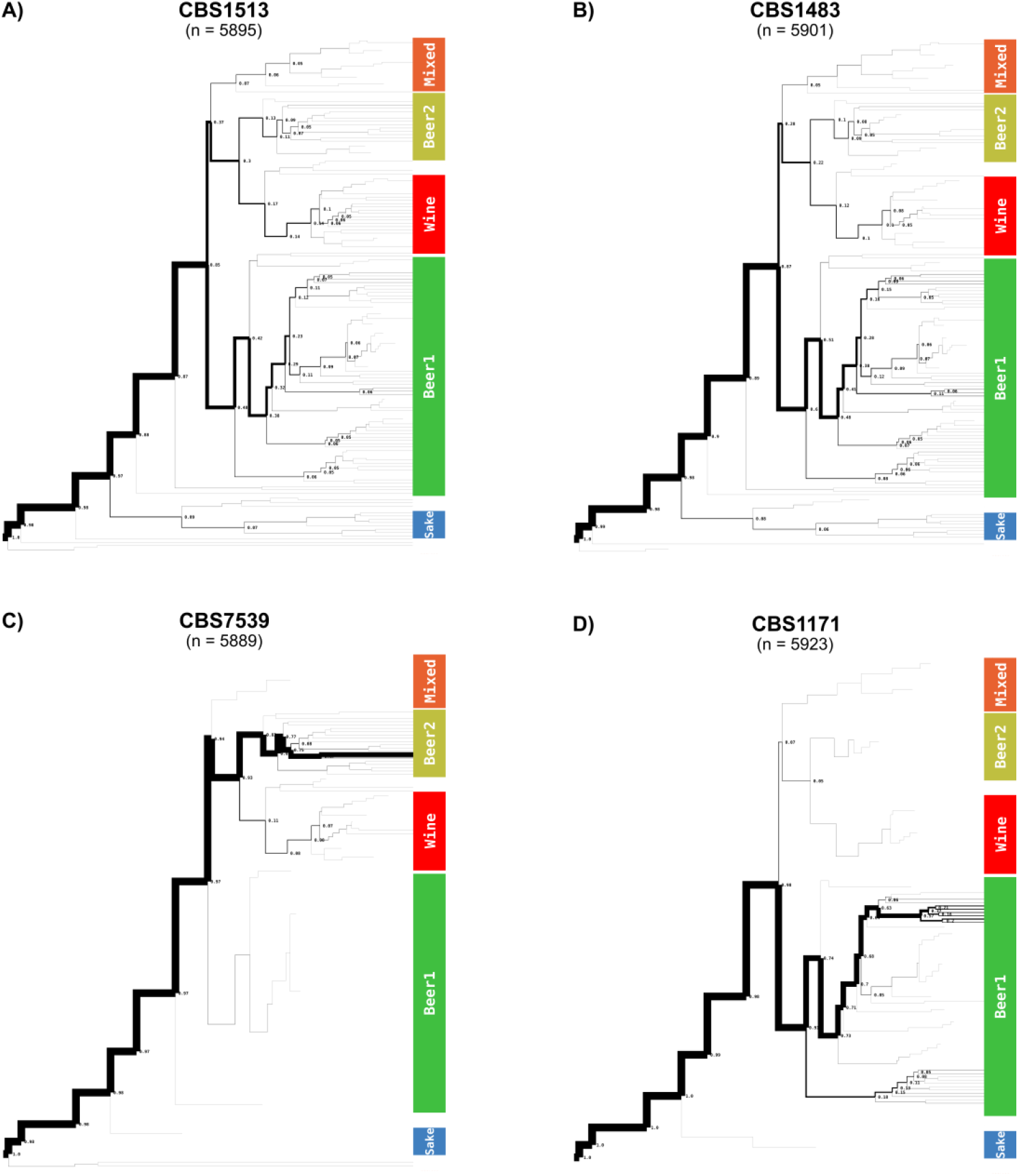
Tree-tracing of the genome-scale similarity across the *S. cerevisiae* (sub-)genomes of various *Saccharomyces* strains, as determined using Alpaca. The frequency at which a genome from the reference data set of 157 *S. cerevisiae* strains from Gallone *et al* (*62*) was identified as most similar for a sub-region of the CBS 1483 genome is depicted. The reference dataset is represented as a population tree, upon which only lineages with similarity are indicated with a thickness proportional to the frequency at which they were found as most similar (‘n’ being the total sum of the number of times all samples appeared as top-scoring). The genomes of *S. pastorianus* Group 1 strain CBS 1513 (A), of *S. pastorianus* Group 2 strain CBS 1483 (B), of *S. cerevisiae* strain CBS 7539 and of *S. cerevisiae* strain CBS 1171 are shown. The tree-tracing figures of *S. pastorianus* Group 1 strains CBS 1503 and CBS 1538, of *S. pastorianus* Group 2 strains CBS 2156, WS34/70 and Hei-A, and of *S. cerevisiae* strains CBS 1463, A81062, CBS 6308, CBS 1487, CEN.PK113-7D and S288C are shown in Figure S4.

To identify possible differences in genome compositions within the *S. cerevisiae* subgenomes of *S. pastorianus*, we analysed other Group 1 and 2 strains using Alpaca, including an isolate of the Heineken A-yeast^®^ lineage (Hei-A), which was isolated in 1886 and represents one of the earliest pure yeast cultures. Whole genome sequencing, alignment to the CBS 1483 assembly and sequencing coverage analysis revealed that the ploidy of the Hei-A isolate corresponds to that of a Group 2 strain (Figure S5). Analysis of Hei-A and the other *S. pastorianus* Group 2 strains CBS 2156 and WS 34/70 using Alpaca yielded almost identical patterns of similarity at the chromosome-level as CBS 1483 (Figure 5). Moreover, similarity was distributed across the *S. cerevisiae* population tree almost identically as in CBS 1483 (Figure 6 and S4). The Group 1 *S. pastorianus* strains CBS 1503, CBS 1513 and CBS 1538 displayed different patterns of similarity at the chromosome-level relative to Group 2 strains. While various chromosome regions harboured almost identical similarity patterns, some regions differed significantly, such as: *Sc*I, the middle of *Sc*IV, the left arm of *Sc*V, *Sc*VIII, the right arm of *Sc*IX, *Sc*X-*Se*X, *Sc*XI and *Sc*XIII (Figure 5). However, at the genome level, similarity was distributed across the *S. cerevisiae* population tree almost identically as in Group 2 strains, except for a slightly higher contribution of the Beer 2 and Wine lineages, at the expense of a lower contribution of the Beer 1 lineage (Figure 6 and S4). The almost identical distribution of all Group 1 and Group 2 strains over the *S. cerevisiae* population tree indicate that they have the same *S. cerevisiae* ancestry. The spread of similarity across the *S. cerevisiae* population tree advocates for an admixed, possibly heterozygous ancestry of the *S. cerevisiae* subgenome of *S. pastorianus*. Furthermore, the different patterns of similarity at the chromosome level between both groups are compatible with an initially heterozygous *S. cerevisiae* subgenome which was subjected to independent loss of heterozygosity events in each group, resulting in differential retention of each haplotype. The lower relative contribution of Beer 1 strains in Group 1 strains may be explained by the complete absence of *S. cerevisiae* chromosomes with high similarity to Beer1 strains, such as ScV, ScXI and *Sc*Xv-*Sc*XI.

## Discussion

In this study, we used Oxford Nanopore Technology’s (ONT) MinION sequencing platform to study the genome of CBS 1483, an alloaneuploid Group 2 *S. pastorianus* strain. The presence of extensively aneuploid *S. cerevisiae* and *S. eubayanus* subgenomes substantially complicates analysis of *S. pastorianus* genomes (10). We therefore explored the ability of nanopore sequencing to generate a reference genome in the presence of multiple non-identical chromosome copies, and investigated the extent to which structural and sequence heterogeneity can be reconstructed. Despite its aneuploidy, we obtained a chromosome-level genome haploid assembly of CBS 1483 in which 29 of the 31 chromosomes were assembled in a single contig. Comparably to assemblies of euploid *Saccharomyces* genomes (22–27), nanopore sequencing resulted in far lesser fragmentation and in the addition of considerable sequences compared to a short-read based assembly of CBS 1483, notably in the subtelomeric regions (10). The added sequences enabled more complete identification of industrially-relevant subtelomeric genes such as the *MAL* genes, responsible for maltose and maltotriose utilisation (37–39), and the *FLO* genes, responsible for flocculation (34, 42, 43). Due to the instability of subtelomeric regions (28–30), the lack of reference-based biases introduced by scaffolding allows more certainty about chromosome structure (24). Since subtelomeric genes encode various industrially-relevant traits (31–34), their mapping enables further progress in strain improvement of lager brewing yeasts. Combined with recently developed Cas9 gene editing tools for *S. pastorianus* (66), accurate localisation and sequence information about subtelomeric genes is critical to investigate their contribution to brewing phenotypes by enabling functional characterization (67).

Despite the presence of non-identical chromosome copies in CBS 1483, the genome assembly only contained one contig per chromosome. While the assembly did not capture information about heterogeneity, mapping of short-read data enabled identification of sequence heterozygosity across the whole genome. In previous work, two alternative chromosome structures could be resolved within a population of euploid *S. cerevisiae* strain CEN.PK113-7D by alignment of nanopore reads (24). Therefore, we evaluated the ability to identify structural heterogeneity by aligning nanopore read data to the assembly. Indeed, nanopore-read alignments enabled the identification of two versions of chromosome *Sc*I: with and without an internal deletion of the gene *UIP3*. Furthermore, the length of nanopore reads enabled them to span a TY-element, revealing that one of the copies of the right arm of *Sc*I was translocated to the left arm of *Sc*XIV. While the two alternative structures of *Sc*I constitute a first step towards the generation of chromosome copy haplotypes, nanopore reads only enabled the hypothesis-based resolution of suspected heterogeneity. Assembly algorithms which do not generate a single consensus sequence per chromosome are emerging (68, 69). However, haplotyping is particularly difficult in aneuploid and polyploid genomes due to copy number differences between chromosomes (68). A further reduction of the relatively high error rate of nanopore reads, or the use of more accurate long-read sequencing technologies, could simplify the generation of haplotype-level genome assemblies in the future by reducing noise (70).

We used the chromosome-level assembly of CBS 1483 to study the ancestry of *S. pastorianus* genomes. Due to the importance of non-linear evolution in the domestication process of *Saccharomyces* strains (56), and to the admixed hybrid nature of *S. pastorianus* (11, 63), we used the newly-developed method Alpaca to analyse the ancestry of CBS 1483 instead of classical phylogenetic approaches using reference datasets of *S. cerevisiae* and *S. eubayanus* strains (62, 63). All *S. pastorianus* genomes displayed identical distribution of similarity across the reference *S. eubayanus* population tree, both at the chromosome and whole-genome level. All *S. pastorianus* genomes also showed identical distribution of similarity across the reference *S. cerevisiae* population tree at the whole genome level; however, Group 1 and Group 2 strains displayed different similarity patterns at the chromosome level. The absence of differences in the *S. cerevisiae* genome at the whole genome level and recurrence of identical chromosomal break points between Group 1 and 2 strains discredit previous hypotheses of different independent hybridization events in the evolution of Group 1 and 2 strains (11, 19). Instead, these results are compatible with the emergence of Group 1 and 2 strains from a single shared hybridization event between a homozygous *S. eubayanus* genome closely related to the Tibetan isolate CDFM21L.1 and an admixed heterozygous *S. cerevisiae* genome with a complex polyphyletic ancestry. Loss of heterozygosity is frequently observed in *Saccharomyces* genomes (56, 71), and therefore likely to have affected both the genomes of Group 1 and 2 strains (11, 72, 73). The different chromosome-level similarity patterns in both groups likely emerged through different loss of heterozygosity events in Group 1 and 2 strains (72, 73). In addition, the lower *S. cerevisiae* chromosome content of Group 1 is consistent with observed loss of genetic material from the least adapted parent during laboratory evolution of *Saccharomyces* hybrids (74–77). In this context, the lower *S. cerevisiae* genome content of Group 1 strains may have resulted from a rare and serendipitous event. For example, chromosome loss has been observed due to unequal chromosome distribution from a sporulation event of a allopolyploid *Saccharomyces* strain (78). Such mutant may have been successful if loss of *S. cerevisiae* chromosomes provided a selective advantage in the low-temperature lager brewing environment (74, 75). The loss of the *S. cerevisiae* subgenome may have affected only Group 1 strains due to different brewing conditions during their domestication. However, the high conservation of similarity within Group 1 and Group 2 strains indicate that the strains within each Group are closely related, indicating a strong population bottleneck in their evolutionary history.

Such a bottleneck could have been caused by the isolation and propagation of a limited number *S. pastorianus* strains, which may have eventually resulted in the extinction of other lineages. The first *S. pastorianus* strains isolated in 1883 by Hansen at the Carlsberg brewery were all Group 1 strains (13, 79). Due to the industry practice of adopting brewing methods and brewing strains from successful breweries, Hansen’s Group 1 isolates likely spread to other breweries as these adopted pure culture brewing (1). Many strains which were identified as Group 2 by whole genome sequencing were isolated in the Netherlands (5, 11): Elion isolated the Heineken A-yeast^®^ in 1886 (80), CBS 1484 was isolated in 1925 from the Oranjeboom brewery (5), CBS 1483 was isolated in 1927 in a Heineken brewery (10), and CBS 1260, CBS 2156 and CBS 5832 were isolated from unknown 16 breweries in the Netherlands in 1937, 1955 and 1968, respectively (5, 81). Analogously to the spread of Group 1 strains from Hansen’s isolate, Group 2 strains may have spread from Elion’s isolate. Both Heineken and Carlsberg distributed their pure culture yeast biomass to breweries over Europe and might therefore have functioned as an evolutionary bottleneck by supplanting other lineages with their isolates (82, 83). Overall, our results support that the differences between Group 1 and 2 strains emerged by differential evolution after an initial shared hybridization event, and not by a different *S. eubayanus* and/or *S. cerevisiae* ancestry.

Beyond its application in this study, we introduced Alpaca as a method to evaluate non-linear evolutionary ancestry. The use of short-read alignments enables Alpaca to account for sequence heterozygosity when assessing similarity between two genomes and is computationally inexpensive as they are reduced to k-mer sets. Moreover, Alpaca leverages previously determined phylogenetic relationships within the reference dataset of strains to infer the evolutionary relationship of the reference genome to the dataset of strains. Due to the presence of non-linear evolutionary processes in a wide range of organisms (84, 85), the applicability of Alpaca extends far beyond the *Saccharomyces* genera. For example, genetic introgressions from *Homo neanderthalensis* constitute about 1% of the human genome (86). Horizontal gene transfer is even relevant across different domains of life: more than 20% of ORFs of the extremely thermophilic bacteria *Thermotoga maritima* were more closely related to genomes of Archaea than to genomes of other Bacteria (87). Critically, horizontal gene transfer, backcrossing and hybridization have not only played a prominent role in the domestication of *Saccharomyces* yeasts (56), but also in other domesticated species such as cows, pigs, wheat and citrus fruits (88–91). Overall, Alpaca can significantly simplify the analysis of new genomes in a broad range of contexts when reference phylogenies are already available.

## Conclusions

With 29 of the 31 chromosomes assembled in single contigs and 323 previously unassembled genes, the genome assembly of CBS 1483 presents the first chromosome-level assembly of a *S. pastorianus* strain specifically, and of an alloaneuploid genome in general. While the assembly only consisted of consensus sequences of all copies of each chromosome, sequence and structural heterozygosity could be recovered by alignment of short and long-reads to the assembly, respectively. We developed Alpaca to investigate the ancestry of Group 1 and Group 2 *S. pastorianus* strains by computing similarity between short-read data from *S. pastorianus* strains relative to large datasets of *S. cerevisiae* and *S. eubayanus* strains. In contrast with the hypothesis of separate hybridization events, Group 1 and 2 strains shared similarity with the same reference *S. cerevisiae* and *S. eubayanus* strains, indicating shared ancestry. Instead, differences between Group 1 and Group 2 strains could be attributed to different patterns of loss of heterozygosity subsequent to a shared hybridization event between a homozygous *S. eubayanus* genome closely related to the Tibetan isolate CDFM21L.1 and an admixed heterozygous *S. cerevisiae* genome with a complex polyphyletic ancestry. We identified the Heineken A-yeast^®^ isolate as a Group 2 strain. We hypothesize that the large differences between Group 1 and Group 2 strains and the high similarity within Group 1 and 2 strains result from a strong population bottleneck which occurred during the isolation of the first Group 1 and Group 2 strains, from which all currently known *S. pastorianus* strains descend. Beyond its application in this study, the ability of Alpaca to reveal non-linear ancestry without requiring heavy computations presents a promising alternative to phylogenetic network analysis to investigate horizontal gene transfer, backcrossing and hybridization.

## Methods

### Yeast strains, cultivation techniques and genomic DNA extraction

*Saccharomyces* strains used in this study are indicated in Table 3. *S. pastorianus* strain CBS 1483, *S. cerevisiae* strain S288C and *S. eubayanus* strain CBS 12357 were obtained from the Westerdijk Fungal Biodiversity Institute (http://www.westerdijkinstitute.nl/). *S. eubayanus* strain CDFM21L.1 was provided by Prof. Feng-Yan Bai. An isolate from the *S. pastorianus* Heineken A-yeast^®^ lineage (Hei-A) was obtained from HEINEKEN Supply Chain B.V., Zoeterwoude, the Netherlands. All strains were stored at −80°C in 30% glycerol (vol/vol). Yeast cultures were inoculated from frozen stocks into 500-mL shake flasks containing 100 mL liquid YPD medium (containing 10 g L^−1^ yeast extract, 20 g L^−1^ peptone and 20 g L^−1^ glucose) and incubated at 12 °C on an orbital shaker set at 200 rpm until the strains reached stationary phase with an OD_660_ between 12 and 20. Genomic DNA was isolated using the Qiagen 100/G kit (Qiagen, Hilden, Germany) according to the manufacturer’s instructions and quantified using a Qubit^®^ Fluorometer 2.0 (ThermoFisher Scientific, Waltham, MA).

**Table 3:** *Saccharomyces* strains used in this study. For strains of the reference dataset, please refer to their original publication (62, 63).

| Name | Species | Descriptio | Reference |
| --- | --- | --- | --- |
| CBS 1483 | *S. pastorianus* | Group 2 | (10) |
| CBS 2156 | *S. pastorianus* | Group 2 | (11) |
| WS 34/70 | *S. pastorianus* | Group 2 | (12) |
| Heineken A-yeast® | *S. pastorianus* | Group 2 | This study |
| CBS 1503 | *S. pastorianus* | Group 1 | (11) |
| CBS 1513 | *S. pastorianus* | Group 1 | (13) |
| CBS 1538 | *S. pastorianus* | Group 1 | (11) |
| S288C | *S. cerevisiae* | Laboratory strain | (102) |
| CEN.PK113-7D | *S. cerevisiae* | Laboratory strain | (24) |
| CBS 7539 | *S. cerevisiae* | Ale brewing strain | (56) |
| CBS 1463 | *S. cerevisiae* | Ale brewing strain | (56) |
| A81062 | *S. cerevisiae* | Ale brewing strain | (18) |
| CBS 1171 | *S. cerevisiae* | Ale brewing strain | (56) |
| CBS 6308 | *S. cerevisiae* | Ale brewing strain | (56) |
| CBS 1487 | *S. cerevisiae* | Ale brewing strain | (56) |
| CBS 12357 | *S. eubayanus* | Patagonian Isolate | (4) |
| CDFM21L.1 | *S. eubayanus* | Tibetan isolate | (65) |

### Short-read Illumina sequencing

Genomic DNA of CBS 1483 and CDFM21L.1 was sequenced on a HiSeq2500 sequencer (Illumina, San Diego, CA) with 125 bp paired-end reads with an insert size of 550 bp using PCR-free library preparation by Keygene (Wageningen, The Netherlands). Genomic DNA of the Heineken A-yeast^®^ isolate Hei-A was sequenced in house on a MiSeq sequencer (Illumina) with 300 bp paired-end reads using PCR-free library preparation. All Illumina sequencing data are available at NCBI (https://www.ncbi.nlm.nih.gov/) under the bioproject accession number PRJNA522669.

### MinION sequencing and basecalling

A total of four MinION genomic libraries of CBS 1483 were created using different chemistries and flow cells: one library using 2D-ligation (Sequencing Kit SQK-MAP006) with a R7.3 chemistry flow cell (FLO-MIN103); two libraries using 2D-ligation (Sequencing Kit SQK-NSK007) with two R9 chemistry flow cells (FLO-MIN105); and one library using 1D-ligation (Sequencing Kit SQK-LASK108) with a R9 chemistry flow cell (FLO-MIN106). All libraries were constructed using the same settings as previously described (24) and reads were uploaded and basecalled using the Metrichor desktop agent (https://metrichor.com/s/). All nanopore sequencing data are available at NCBI (https://www.ncbi.nlm.nih.gov/) under the bioproject accession number PRJNA522669.

### *De novo* genome assembly

The genome of CBS 1483 was assembled *de novo* using only the ONT sequencing data generated in this study. The assembly was generated using Canu (35), polished using Pilon (36) and annotated using MAKER2 (92), as previously described (24) with some modifications: Pilon (version 1.22) was only used to polish sequencing errors in the ONT-only *de* novo assembly, and Minimap2 (93) (version 2.7) was used as the long-read aligner to identify potential misassemblies and heterozygous structural variants, which were visualized using Ribbon (94). The resulting assembly was manually curated: (i) a contig of 24 Kbp comprised entirely of “TATATA” sequence was discarded; (ii) three contigs of 592, 465, and 95 Kbp (corresponding to the rDNA locus of the *S. cerevisiae* sub-genome) and complete sequence up and downstream of this locus were joined with a gap; (iii) four contigs corresponding to *S. cerevisiae* chromosome I (referred to as *Sc*I) were joined without a gap into a complete 208 Kbp chromosome assembly (Figure 2A); (iv) two contigs corresponding to *Sc*XIV were joined with a gap (Figure 2D); and (v) 23 Kbp of overlapping sequence from the mitochondrial contig corresponding to the origin of replication was identified with Nucmer (95) and manually removed when circularizing the contig, leading to the complete a final size of 69 Kbp. The assembled genomes are available at NCBI (https://www.ncbi.nlm.nih.gov/) under the bioproject accession number PRJNA522669. Gene annotations are available in Additional File 1A.

### Comparison between ONT-only and Illumina-only genome assembly

Gained and lost sequence information in the nanopore assembly of CBS 1483 was determined by comparing it to the previous short-read assembly (10), as previously described (24) with the addition of using minimum added sequence length of 25 nt.

### FLO gene analysis

We used Tandem Repeat Finder (version 4.09) (96) with recommended parameters to identify tandem repeat sequences in *FLO1* (SGDID:S000000084), *FLO5* (SGDID:S000001254), *FLO8* (SGDID:S000000911), *FLO9* (SGDID:S000000059), *FLO10* (SGDID:S000001810), and *FLO11* (SGDID:S000001458) of *S. cerevisiae* strain S288C (97) as well as in *FLO1*, *FLO5*, *FLO8*, *FLO9*, *FLO10* and *FLO11* of *S. eubayanus* strain CBS 12357 (25). The resulting tandem repeat sequences were then used as proxies to characterize *FLO* genes in our assembly of CBS 1483, in a previously generated assembly of *S. cerevisiae* strain CEN.PK113-7D (24) and the *Lg-FLO1* genes previously described in *S. cerevisiae* strain CMBSVM11 (GenBank HM358276) and *S. pastorianus* strain KBY001 (GenBank D89860.1) (51, 52). BLASTN (version 2.2.31+) (98) was then used to align the tandem sequences to each FLO gene. The alignments were further processed via an in-house script in the Scala programming language to identify repeat clusters by requiring a minimum alignment coverage of 0.5 and a maximum gap between two repeats of 3x times the repeat sequence length. The total number of copies was estimated by dividing the total size of the cluster by the repeat sequence length.

### Intra-chromosomal heterozygosity

Sequence variation was identified by aligning the short-read Illumina reads generated in this study to the ONT-only assembly with BWA (99) and calling variants with Pilon (36) using the *--fix “bases”*, *”local”* and *--diploid* parameters. To restrict false positive calls, SNPs were disregarded within 10 Kbp of the ends of the chromosomes, if minor alleles had a frequency below 15% allele frequency, and if the coverage was below 3 reads.

Copy number variation for all chromosomes were estimated by aligning all short-reads to the ONT-only assembly. Reads were trimmed of adapter sequences and low-quality bases with Trimmomatic (100) (version 0.36) and aligned with BWA (99) (version 0.7.12). The median coverage was computed using a non-overlapping window of 100 nt, copy number was determined by comparing the coverage to that of the chromosome with the smallest median coverage. Additionally, copy number variation at the gene-level was also investigated based on whether the coverage of an individual gene significantly deviated from the coverage of the surrounding region. First, we defined contiguous chromosomal sub-regions with fixed copy number (Table S1). The mean and standard deviation of coverages of these sub-regions were then computed using ONT-only alignments. Mean coverages of every gene was then computed and an uncorrected Z-test (101) was performed by comparing a gene’s mean coverage and the corresponding mean and standard deviation of the pre-defined sub-region that the gene overlapped with.

### Similarity analysis and lineage tracing of *S. pastorianus* sub-genomes using Alpaca

We developed Alpaca (61) to investigate non-linear ancestry of a reference genome based on large sequencing datasets. Briefly, Alpaca partitions a reference genome into multiple sub-regions, each reduced to a k-mer set representation. Sequence similarities of the sub-regions are then independently computed against the corresponding sub-regions in a collection of target genomes. Non-linear ancestry can therefore be inferred by tracing the population origin of the most similar genome(s) in each sub-region. Detailed explanation Alpaca can be found in our method description (61).

Alpaca (version 1.0) was applied to the ONT CBS 1483 genome assembly to investigate the similarity of sub-regions from both sub-genomes to previously defined population lineages. For partitioning the CBS 1483 genome into sub-regions, we used a k-mer size of 21 and a sub-region size of 2 Kbp and used the short-read Illumina data of CBS 1483 produced in this study to assure accurate k-mer set construction. For investigating mosaic structures in the *S. cerevisiae* subgenome, we used 157 brewing-related *S. cerevisiae* genomes (project accession number PRJNA323691) which were subdivided in six major lineages: Asia, Beer1, Beer2, Mixed, West-Africa, Wine and Mosaic (62). For the *S. eubayanus* subgenome, we used 29 available genomes (project accession number PRJNA290017) which were subdivided in three major lineages: Admixed, Patagonia-A, and Patagonia-B (63). Raw-reads of all samples were trimmed Trimmomatic and filtered reads were aligned to CBS 1483 genome using BWA (99). Alpaca was also applied to several *Saccharomyces* genomes to investigate evolutionary similarities and differences between Group 1 and Group 2 *S. pastorianus* genomes. We used Group 1 strains CBS 1503, CBS 1513, and CBS 1538, and Group 2 strains CBS 2156 and WS34/70 (project accession number PRJDB4073) (11). As a control, eight *S. cerevisiae* genomes were analysed: ale strains CBS 7539, CBS 1463, CBS 1171, CBS 6308, and CBS 1487 (project accession number PRJEB13017) (56) and A81062 (project accession number PRJNA408119) (18), and laboratory strains CEN.PK113-7D (project accession number PRJNA393501) (24) and S288C (project accession number PRJEB14774) (23). Similarly, raw-reads for all strains were trimmed with Trimmomatic and aligned to the ONT CBS 1483 genome assembly using BWA. Partitioning of the additional *S. pastorianus* and *S. cerevisiae* genomes with Alpaca was performed by deriving k-mer sets from read-alignments only, assuring direct one-to-one comparison of all sub-regions across all genomes. K-mer size of 21 and sub-region size of 2 Kbp were used. The *S. cerevisiae* and *S. eubayanus* sequencing data were used to identify potential mosaic structures in these genomes. Lastly, *S. cerevisiae* and *S. eubayanus* strains were subdivided into subpopulations according to previously defined lineages (62, 63). MASH (version 2.1) (*64*) was then used to hierarchically cluster each genome based on their MASH distance using k-mer size of 21, sketch size of 1,000,000, and minimum k-mer frequency of 2. The resulting trees were used as population reference trees for Alpaca (61).

## Declarations

### Ethics approval and consent to participate

Not applicable

### Consent for publication

Not applicable

### Availability of data and materials

The datasets generated and/or analysed during the current study are available in the NCBI repository, https://www.ncbi.nlm.nih.gov/.

### Authors’ contributions

ANS and PdlTC performed nanopore sequencing. NB performed illumina sequencing. ANS performed sequence assembly. ANS, ARGDV and MvdB analysed the data. ANS designed and applied Alpaca. HEINEKEN Supply Chain B.V. provided the Heineken A-yeast^®^ isolate. NGA supported sequencing and reviewed the manuscript. ANS and ARGDV wrote the manuscript. JMGD and TA supervised the study. All authors read and approved the final manuscript.

## Acknowledgments

We thank Prof. Feng Yan Bai for kindly providing us *S. eubayanus* strain CDFM21L.1, as well as Prof. Jack Pronk and Dr. Jan-Maarten Geertman for their support throughout the study.

## Supplemental file, figure and table legends

**Additional File 1A:** Gene annotations of the nanopore assembly of CBS 1483 as predicted by MAKER2.

**Additional File 1B:** Added sequence in the nanopore assembly relative to the van den Broek et al 2015 assembly.

**Additional File 1C:** Lost sequence in the nanopore assembly relative to the van den Broek et al 2015 assembly.

**Additional File 1D:** Gene ontology analysis of genes identified in the nanopore assembly which were absent in the van den Broek et al 2015 assembly.

**Additional file 1E:** Genes from brewing-relevant subtelomeric gene families in the nanopore assembly of CBS 1483

**Additional file 1F:** Heterozygous sequences in the nanopore assembly of CBS 1483

**Additional file 1G:** ORFs with copy number deviating from the copy number of surrounding sequences in the nanopore assembly of CBS 1483

**Additional File 1H:** Sequences of the repeats identified in FLO genes of *S. cerevisiae* S288C and *S. eubayanus* CBS 12357.

**Figure S1.** Read-length distribution obtained of the MinION libraries of CBS 1483 produced in this study. This plot shows the four different sequencing libraries obtained from whole genome sequencing of CBS 1483 using the MinION^®^ platform. The Y-axes the frequency and the X-axes depict read-lengths (capped at 30 Kbp) using bins of 500 bp. The libraries were obtained with different sequencing chemistries due to the rapid development of Nanopore sequencing.

**Figure S2.** The ploidy estimates for each chromosome in the CBS 1483 assembly based on Illumina short read data from this study. The red horizontal lines correspond to the median coverage of each chromosome.

**Figure S3.** Copy number predictions for ORFs in the genome assembly of CBS 1483. Each dot represents an ORF whose x-value represents the location in the corresponding chromosome and the y-value the estimated copy number based on coverage analysis of ONT-only read alignments. Dots in yellow indicate ORFs whose coverage significantly deviates from the surrounding genomic region, indicating copy number variation of the ORF within different copies of the same chromosome.

**Figure S4.** More complete version of the tree tracing of Figure 6. Tree-tracing of the genome-scale similarity across the *S. cerevisiae* (sub-)genomes of various *Saccharomyces* strains, as determined using Alpaca. The frequency at which a genome from the reference data set of 157 *S. cerevisiae* strains from Gallone *et al* (*62*) was identified as most similar for a sub-region of the CBS 1483 genome is depicted. The reference dataset is represented as a population tree, upon which only lineages with similarity are indicated with a thickness proportional to the frequency at which they were found as most similar. In addition to the genomes of CBS 1513, CBS 1483, CBS 7539 and CBS 1171, the tree-tracing figures of *S. pastorianus* Group 1 strains CBS 1503 and CBS 1538, of *S*. *pastorianus* Group 2 strains CBS 2156, WS34/70 and Hei-A, and of *S. cerevisiae* strains CBS 1463, A81062, CBS 6308, CBS 1487, CEN.PK113-7D and S288C are shown.

**Figure S5.** The ploidy estimates for each chromosome of the *S. pastorianus* isolate of the Heineken A-yeast^®^ lineage based on alignment of short-read data to the chromosome-level assembly of CBS 1483.

**Table S1.** Contiguous sub-regions in the CBS 1483 genome with fixed copy number.

## References

1. Meussdoerffer FG. A comprehensive history of beer brewing. In: Eßlinger HM, editor. Handbook of brewing: Processes, technology, markets. Weinheim: Wiley-VCH; 2009. p. 1–42.

2. Kodama Y, Kielland-Brandt MC, Hansen J. Lager brewing yeast. Comparative genomics. Berlin: Springer; 2006. p. 145–64.

3. Dequin S. The potential of genetic engineering for improving brewing, wine-making and baking yeasts. Appl Microbiol Biotechnol. 2001;56(5-6):577–88.

4. Libkind D, Hittinger CT, Valério E, Gonçalves C, Dover J, Johnston M, et al. Microbe domestication and the identification of the wild genetic stock of lager-brewing yeast. Proc Natl Acad Sci U S A. 2011;108(35):14539–44.

5. Dunn B, Sherlock G. Reconstruction of the genome origins and evolution of the hybrid lager yeast *Saccharomyces pastorianus*. Genome Res. 2008;18(10):1610–23.

6. de Barros Lopes M, Bellon JR, Shirley NJ, Ganter PF. Evidence for multiple interspecific hybridization in *Saccharomyces* sensu stricto species. FEMS Yeast Res. 2002;1(4):323–31.

7. Hebly M, Brickwedde A, Bolat I, Driessen MR, de Hulster EA, van den Broek M, et al. *S. cerevisiae* × *S. eubayanus* interspecific hybrid, the best of both worlds and beyond. FEMS Yeast Res. 2015;15(3).

8. Krogerus K, Magalhães F, Vidgren V, Gibson B. New lager yeast strains generated by interspecific hybridization. J Ind Microbiol Biotechnol. 2015;42(5):769–78.

9. Gorter de Vries AR, Pronk JT, Daran J-MG. Industrial relevance of chromosomal copy number variation in *Saccharomyces* yeasts. Appl Environ Microbiol. 2017:AEM. 03206–16.

10. Van den Broek M, Bolat I, Nijkamp J, Ramos E, Luttik MA, Koopman F, et al. Chromosomal copy number variation in *Saccharomyces pastorianus* evidence for extensive genome dynamics in industrial lager brewing strains. Appl Environ Microbiol. 2015:AEM. 01263–15.

11. Okuno M, Kajitani R, Ryusui R, Morimoto H, Kodama Y, Itoh T. Next-generation sequencing analysis of lager brewing yeast strains reveals the evolutionary history of interspecies hybridization. DNA Res. 2016;23(1):67–80.

12. Nakao Y, Kanamori T, Itoh T, Kodama Y, Rainieri S, Nakamura N, et al. Genome sequence of the lager brewing yeast, an interspecies hybrid. DNA Res. 2009;16(2):115–29.

13. Walther A, Hesselbart A, Wendland J. Genome sequence of *Saccharomyces carlsbergensis*, the world’s first pure culture lager yeast. G3 (Bethesda). 2014:g3. 113.010090.

14. Hewitt SK, Donaldson IJ, Lovell SC, Delneri D. Sequencing and characterisation of rearrangements in three *S. pastorianus* strains reveals the presence of chimeric genes and gives evidence of breakpoint reuse. PLoS One. 2014;9(3):e92203.

15. Liti G, Peruffo A, James SA, Roberts IN, Louis EJ. Inferences of evolutionary relationships from a population survey of LTR-retrotransposons and telomeric-associated sequences in the *Saccharomyces* sensu stricto complex. Yeast. 2005;22(3):177–92.

16. Monerawela C, James TC, Wolfe KH, Bond U. Loss of lager specific genes and subtelomeric regions define two different *Saccharomyces cerevisiae* lineages for *Saccharomyces pastorianus* Group I and II strains. FEMS Yeast Res. 2015;15(2):fou008.

17. Rainieri S, Kodama Y, Kaneko Y, Mikata K, Nakao Y, Ashikari T. Pure and mixed genetic lines of *Saccharomyces bayanus* and *Saccharomyces pastorianus* and their contribution to the lager brewing strain genome. Appl Environ Microbiol. 2006;72(6):3968–74.

18. Krogerus K, Arvas M, De Chiara M, Magalhães F, Mattinen L, Oja M, et al. Ploidy influences the functional attributes of *de novo* lager yeast hybrids. Appl Microbiol Biotechnol. 2016;100(16):7203–22.

19. Baker E, Wang B, Bellora N, Peris D, Hulfachor AB, Koshalek JA, et al. The genome sequence of *Saccharomyces eubayanus* and the domestication of lager-brewing yeasts. Mol Biol Evol. 2015;32(11):2818–31.

20. Gorter de Vries A, Voskamp MA, van Aalst ACA, Kristensen LH, Jansen L, van den Broek M, et al. Laboratory evolution of a *Saccharomyces cerevisiae* × *S. eubayanus* hybrid under simulated lager-brewing conditions: genetic diversity and phenotypic convergence. bioRxiv. 2018.

21. Kim JM, Vanguri S, Boeke JD, Gabriel A, Voytas DF. Transposable elements and genome organization: a comprehensive survey of retrotransposons revealed by the complete *Saccharomyces cerevisiae* genome sequence. Genome Res. 1998;8(5):464–78.

22. Giordano F, Aigrain L, Quail MA, Coupland P, Bonfield JK, Davies RM, et al. *De novo* yeast genome assemblies from MinION, PacBio and MiSeq platforms. Sci Rep. 2017;7(1):3935.

23. Istace B, Friedrich A, d’Agata L, Faye S, Payen E, Beluche O, et al. *de novo* assembly and population genomic survey of natural yeast isolates with the Oxford Nanopore MinION sequencer. Gigascience. 2017;6(2):1–13.

24. Salazar AN, Gorter de Vries AR, van den Broek M, Wijsman M, de la Torre Cortés P, Brickwedde A, et al. Nanopore sequencing enables near-complete *de novo* assembly of *Saccharomyces cerevisiae* reference strain CEN. PK113-7D. FEMS Yeast Res. 2017;17(7).

25. Brickwedde A, Brouwers N, van den Broek M, Gallego Murillo JS, Fraiture JL, Pronk JT, et al. Structural, physiological and regulatory analysis of maltose transporter genes in *Saccharomyces eubayanus* CBS 12357T. Front Microbiol. 2018;9:1786.

26. Yue J-X, Li J, Aigrain L, Hallin J, Persson K, Oliver K, et al. Contrasting evolutionary genome dynamics between domesticated and wild yeasts. Nat Genet. 2017;49(6):913.

27. McIlwain SJ, Peris D, Sardi M, Moskvin OV, Zhan F, Myers K, et al. Genome sequence and analysis of a stress-tolerant, wild-derived strain of *Saccharomyces cerevisiae* used in biofuels research. G3 (Bethesda). 2016:g3. 116.029389.

28. Pryde FE, Huckle TC, Louis EJ. Sequence analysis of the right end of chromosome XV in *Saccharomyces cerevisiae*: an insight into the structural and functional significance of sub-telomeric repeat sequences. Yeast. 1995;11(4):371–82.

29. Bergström A, Simpson JT, Salinas F, Barré B, Parts L, Zia A, et al. A high-definition view of functional genetic variation from natural yeast genomes. Mol Biol Evol. 2014;31(4):872–88.

30. Brown CA, Murray AW, Verstrepen KJ. Rapid expansion and functional divergence of subtelomeric gene families in yeasts. Curr Biol. 2010;20(10):895–903.

31. Jordan P, Choe J-Y, Boles E, Oreb M. Hxt13, Hxt15, Hxt16 and Hxt17 from *Saccharomyces cerevisiae* represent a novel type of polyol transporters. Sci Rep. 2016;6:23502.

32. Teste M-A, François JM, Parrou J-L. Characterization of a new multigene family encoding isomaltases in the yeast *Saccharomyces Cerevisiae*: the *IMA* family. J Biol Chem. 2010:jbc. M110. 145946.

33. Denayrolles M, de Villechenon EP, Lonvaud-Funel A, Aigle M. Incidence of *SUC-RTM* telomeric repeated genes in brewing and wild wine strains of *Saccharomyces*. Curr Genet. 1997;31(6):457–61.

34. Teunissen A, Steensma H. The dominant flocculation genes of Saccharomyces cerevisiae constitute a new subtelomeric gene family. Yeast. 1995;11(11):1001–13.

35. Koren S, Walenz BP, Berlin K, Miller JR, Bergman NH, Phillippy AM. Canu: scalable and accurate long-read assembly via adaptive k-mer weighting and repeat separation. Genome Res. 2017:gr. 215087.116.

36. Walker BJ, Abeel T, Shea T, Priest M, Abouelliel A, Sakthikumar S, et al. Pilon: an integrated tool for comprehensive microbial variant detection and genome assembly improvement. PLoS One. 2014;9(11):e112963.

37. Alves SL, Herberts RA, Hollatz C, Trichez D, Miletti LC, De Araujo PS, et al. Molecular analysis of maltotriose active transport and fermentation by *Saccharomyces cerevisiae* reveals a determinant role for the *AGT1* permease. Appl Environ Microbiol. 2008;74(5):1494–501.

38. Chang Y, Dubin R, Perkins E, Michels C, Needleman R. Identification and characterization of the maltose permease in genetically defined *Saccharomyces* strain. J Bacteriol. 1989;171(11):6148–54.

39. Naumov GI, Naumova ES, Michels C. Genetic variation of the repeated *MAL* loci in natural populations of *Saccharomyces cerevisiae* and *Saccharomyces paradoxus*. Genetics. 1994;136(3):803–12.

40. Zastrow C, Hollatz C, De Araujo P, Stambuk B. Maltotriose fermentation by *Saccharomyces cerevisiae*. J Ind Microbiol Biotechnol. 2001;27(1):34–8.

41. Baker EP, Hittinger CT. Evolution of a novel chimeric maltotriose transporter in *Saccharomyces eubayanus* from parent proteins unable to perform this function. bioRxiv. 2018.

42. Van Mulders SE, Christianen E, Saerens SM, Daenen L, Verbelen PJ, Willaert R, et al. Phenotypic diversity of Flo protein family-mediated adhesion in *Saccharomyces cerevisiae*. FEMS Yeast Res. 2009;9(2):178–90.

43. Miki B, Poon NH, James AP, Seligy VL. Possible mechanism for flocculation interactions governed by gene *FLO1* in *Saccharomyces cerevisiae*. J Bacteriol. 1982;150(2):878–89.

44. Dengis PB, Nelissen L, Rouxhet PG. Mechanisms of yeast flocculation: comparison of top- and bottom-fermenting strains. Appl Environ Microbiol. 1995;61(2):718–28.

45. Fidalgo M, Barrales RR, Jimenez J. Coding repeat instability in the *FLO11* gene of *Saccharomyces yeasts*. Yeast. 2008;25(12):879–89.

46. Zara G, Zara S, Pinna C, Marceddu S, Budroni M. FLO11 gene length and transcriptional level affect biofilm-forming ability of wild flor strains of *Saccharomyces cerevisiae*. Microbiology. 2009;155(12):3838–46.

47. Verstrepen KJ, Jansen A, Lewitter F, Fink GR. Intragenic tandem repeats generate functional variability. Nat Genet. 2005;37(9):986.

48. Liu N, Wang D, Wang ZY, He XP, Zhang B. Genetic basis of flocculation phenotype conversion in *Saccharomyces cerevisiae*. FEMS Yeast Res. 2007;7(8):1362–70.

49. Ogata T, Izumikawa M, Kohno K, Shibata K. Chromosomal location of Lg-*FLO1* in bottom-fermenting yeast and the *FLO5* locus of industrial yeast. J Appl Microbiol. 2008;105(4):1186–98.

50. Soares EV. Flocculation in *Saccharomyces cerevisiae*: a review. J Appl Microbiol. 2011;110(1):1–18.

51. Van Mulders SE, Ghequire M, Daenen L, Verbelen PJ, Verstrepen KJ, Delvaux FR. Flocculation gene variability in industrial brewer’s yeast strains. Appl Microbiol Biotechnol. 2010;88(6):1321–31.

52. Kobayashi O, Hayashi N, Kuroki R, Sone H. Region of Flo1 proteins responsible for sugar recognition. J Bacteriol. 1998;180(24):6503–10.

53. Li P, Guo X, Shi T, Hu Z, Chen Y, Du L, et al. Reducing diacetyl production of wine by overexpressing *BDH1* and *BDH2* in *Saccharomyces uvarum*. J Ind Microbiol Biotechnol. 2017;44(11):1541–50.

54. Saerens S, Thevelein J, Delvaux F. Ethyl ester production during brewery fermentation, a review. Cerevisia. 2008;33(2):82–90.

55. Gibson BR, Storgårds E, Krogerus K, Vidgren V. Comparative physiology and fermentation performance of Saaz and Frohberg lager yeast strains and the parental species *Saccharomyces eubayanus*. Yeast. 2013;30(7):255–66.

56. Peter J, De Chiara M, Friedrich A, Yue J-X, Pflieger D, Bergström A, et al. Genome evolution across 1,011 *Saccharomyces cerevisiae* isolates. Nature. 2018;556(7701):339.

57. Gogarten JP, Townsend JP. Horizontal gene transfer, genome innovation and evolution. Nat Rev Microbiol. 2005;3(9):679.

58. Solís-Lemus C, Bastide P, Ané C. PhyloNetworks: a package for phylogenetic networks. Mol Biol Evol. 2017;34(12):3292–8.

59. Hejase HA, Liu KJ. A scalability study of phylogenetic network inference methods using empirical datasets and simulations involving a single reticulation. BMC bioinformatics. 2016;17(1):422.

60. Consortium CP-G. Computational pan-genomics: status, promises and challenges. Brief Bioinform. 2016;19(1):118–35.

61. Salazar A, Abeel T. Alpaca: a kmer-based approach for investigating mosaic structures in microbial genomes. bioRxiv. 2019:551234.

62. Gallone B, Steensels J, Prahl T, Soriaga L, Saels V, Herrera-Malaver B, et al. Domestication and divergence of *Saccharomyces cerevisiae* beer yeasts. Cell. 2016;166(6):1397–410. e16.

63. Peris D, Langdon QK, Moriarty RV, Sylvester K, Bontrager M, Charron G, et al. Complex ancestries of lager-brewing hybrids were shaped by standing variation in the wild yeast *Saccharomyces eubayanus*. PLoS Genet. 2016;12(7):e1006155.

64. Ondov BD, Treangen TJ, Melsted P, Mallonee AB, Bergman NH, Koren S, et al. Mash: fast genome and metagenome distance estimation using MinHash. Genome Biol. 2016;17(1):132.

65. Bing J, Han P-J, Liu W-Q, Wang Q-M, Bai F-Y. Evidence for a Far East Asian origin of lager beer yeast. Curr Biol. 2014;24(10):R380–R1.

66. Gorter de Vries AR, Groot PA, Broek M, Daran J-MG. CRISPR-Cas9 mediated gene deletions in lager yeast *Saccharomyces pastorianus*. Microb Cell Fact. 2017;16(1):222.

67. Winzeler EA, Shoemaker DD, Astromoff A, Liang H, Anderson K, Andre B, et al. Functional characterization of the *S. cerevisiae* genome by gene deletion and parallel analysis. Science. 1999;285(5429):901–6.

68. He D, Saha S, Finkers R, Parida L. Efficient algorithms for polyploid haplotype phasing. BMC genomics. 2018;19(2):110.

69. Chin C-S, Peluso P, Sedlazeck FJ, Nattestad M, Concepcion GT, Clum A, et al. Phased diploid genome assembly with single-molecule real-time sequencing. Nat Methods. 2016;13(12):1050.

70. Wenger AM, Peluso P, Rowell WJ, Chang P-C, Hall RJ, Concepcion GT, et al. Highly-accurate long-read sequencing improves variant detection and assembly of a human genome. bioRxiv. 2019:519025.

71. Magwene PM, Kayıkçı Ö, Granek JA, Reininga JM, Scholl Z, Murray D. Outcrossing, mitotic recombination, and life-history trade-offs shape genome evolution in *Saccharomyces cerevisiae*. Proc Natl Acad Sci U S A. 2011;108(5):1987–92.

72. Chambers SR, Hunter N, Louis EJ, Borts RH. The mismatch repair system reduces meiotic homeologous recombination and stimulates recombination-dependent chromosome loss. Mol Cell Biol. 1996;16(11):6110–20.

73. González SS, Barrio E, Querol A. Molecular characterization of new natural hybrids of *Saccharomyces cerevisiae* and *S. kudriavzevii* in brewing. Appl Environ Microbiol. 2008;74(8):2314–20.

74. Smukowski Heil CS, DeSevo CG, Pai DA, Tucker CM, Hoang ML, Dunham MJ. Loss of heterozygosity drives adaptation in hybrid yeast. Mol Biol Evol. 2017;34(7):1596–612.

75. Heil CS, Large CR, Patterson K, Dunham MJ. Temperature preference biases parental genome retention during hybrid evolution. bioRxiv. 2018:429803.

76. Pérez Través L, Lopes CA, Barrio E, Querol A. Study of the stabilization process in *Saccharomyces* intra- and interspecific hybrids in fermentation conditions. Int Microbiol. 2014;17(4):213–24.

77. Antunovics Z, Nguyen H-V, Gaillardin C, Sipiczki M. Gradual genome stabilisation by progressive reduction of the *Saccharomyces uvarum* genome in an interspecific hybrid with *Saccharomyces cerevisiae*. FEMS Yeast Res. 2005;5(12):1141–50.

78. Lopandic K, Pfliegler WP, Tiefenbrunner W, Gangl H, Sipiczki M, Sterflinger K. Genotypic and phenotypic evolution of yeast interspecies hybrids during high-sugar fermentation. Appl Microbiol Biotechnol. 2016;100(14):6331–43.

79. Hansen EC. Recherches sur la physiologie et la morphologie des ferments alcooliques. V. Methodes pour obtenir des cultures pures de Saccharomyces et de microorganismes analogues. Compt Rend Trav Lab Carlsberg. 1883;2:92–105.

80. Gélinas P. Mapping early patents on baker’s yeast manufacture. Compr Rev Food Sci Food Saf. 2010;9(5):483–97.

81. Scheda R, Yarrow D. Variation in the fermentative pattern of some *Saccharomyces* species. Arch Mikrobiol. 1968;61(3):310–6.

82. Hornsey IS. A history of beer and brewing. Cambridge, UK: Royal Society of Chemistry; 2003.

83. Mendlik F. Some aspects of the scientific development of brewing in Holland. J Inst Brew. 1937;43(4):294–300.

84. Keeling PJ, Palmer JD. Horizontal gene transfer in eukaryotic evolution. Nat Rev Genet. 2008;9(8):605.

85. Thomas CM, Nielsen KM. Mechanisms of, and barriers to, horizontal gene transfer between bacteria. Nat Rev Microbiol. 2005;3(9):711.

86. Racimo F, Sankararaman S, Nielsen R, Huerta-Sánchez E. Evidence for archaic adaptive introgression in humans. Nat Rev Genet. 2015;16(6):359.

87. Nelson KE, Clayton RA, Gill SR, Gwinn ML, Dodson RJ, Haft DH, et al. Evidence for lateral gene transfer between Archaea and bacteria from genome sequence of *Thermotoga maritima*. Nature. 1999;399(6734):323.

88. Larson G, Dobney K, Albarella U, Fang M, Matisoo-Smith E, Robins J, et al. Worldwide phylogeography of wild boar reveals multiple centers of pig domestication. Science. 2005;307(5715):1618–21.

89. McTavish EJ, Decker JE, Schnabel RD, Taylor JF, Hillis DM. New World cattle show ancestry from multiple independent domestication events. Proc Natl Acad Sci U S A. 2013;110(15):E1398–E406.

90. Brenchley R, Spannagl M, Pfeifer M, Barker GL, D’Amore R, Allen AM, et al. Analysis of the bread wheat genome using whole-genome shotgun sequencing. Nature. 2012;491(7426):705.

91. Wu GA, Prochnik S, Jenkins J, Salse J, Hellsten U, Murat F, et al. Sequencing of diverse mandarin, pummelo and orange genomes reveals complex history of admixture during citrus domestication. Nat Biotechnol. 2014;32(7):656.

92. Holt C, Yandell M. MAKER2: an annotation pipeline and genome-database management tool for second-generation genome projects. BMC bioinformatics. 2011;12(1):491.

93. Li H. Minimap2: pairwise alignment for nucleotide sequences. Bioinformatics. 2018;34(18):3094–100.

94. Nattestad M, Chin C-S, Schatz MC. Ribbon: visualizing complex genome alignments and structural variation. bioRxiv. 2016:082123.

95. Kurtz S, Phillippy A, Delcher AL, Smoot M, Shumway M, Antonescu C, et al. Versatile and open software for comparing large genomes. Genome Biol. 2004;5(2):R12.

96. Benson G. Tandem repeats finder: a program to analyze DNA sequences. Nucleic acids research. 1999;27(2):573–80.

97. Cherry JM, Hong EL, Amundsen C, Balakrishnan R, Binkley G, Chan ET, et al. Saccharomyces Genome Database: the genomics resource of budding yeast. Nucleic acids research. 2011;40(D1):D700–D5.

98. Camacho C, Coulouris G, Avagyan V, Ma N, Papadopoulos J, Bealer K, et al. BLAST+: architecture and applications. BMC bioinformatics. 2009;10(1):421.

99. Li H, Durbin R. Fast and accurate long-read alignment with Burrows–Wheeler transform. Bioinformatics. 2010;26(5):589–95.

100. Bolger AM, Lohse M, Usadel B. Trimmomatic: a flexible trimmer for Illumina sequence data. Bioinformatics. 2014;30(15):2114–20.

101. Fisher RA. The design of experiments: Oliver And Boyd; Edinburgh; London; 1937.

102. Goffeau A, Barrell BG, Bussey H, Davis R, Dujon B, Feldmann H, et al. Life with 6000 genes. Science. 1996;274(5287):546–67.

